# Mechanistic Study of palmatine in Regulating Pyroptosis in Sepsis Cells via Signaling Pathways

**DOI:** 10.1101/2025.02.15.638474

**Authors:** Guangzhao Yan, Yanyan Wang

## Abstract

This study aims to investigate whether palmatine has a potential effect on sepsis-related pyroptosis in patients through relevant signaling pathways. Initially, the study explores the differential genes associated with pyroptosis in sepsis patients, along with their related enrichment processes, biological behaviors, and functional pathways. Patients with sepsis pyroptosis are then classified into subgroups based on pyroptosis-related genes to analyze the biological differences between these subtypes. Furthermore, the signaling pathways and biological functions associated with palmatine are studied to assess whether there are functional similarities or overlaps with the signaling pathways and biological behaviors observed in patients with sepsis pyroptosis. Finally, the findings are validated through wet-lab experiments. The results reveal that the signaling pathways related to palmatine partially overlap with those observed in sepsis pyroptosis patients, indicating that palmatine may influence sepsis pyroptosis through these pathways. Parmatine may improve the prognosis of sepsis by affecting the immune function of PRKACA, PTGS2, NLRP3, HSP90AA1 and PTPN22.

## Introduction

Sepsis is a common critical illness characterized by life-threatening organ dysfunction caused by an abnormal bodily response to infection. The development of sepsis involves a variety of complex pathophysiological processes, including cellular dysfunction, uncontrolled inflammatory responses, microcirculatory failure, and coagulation abnormalities [1,2]. According to statistics [3], the global incidence of sepsis reaches millions of cases annually, with a particularly high burden in low- and middle-income countries, where sepsis patients face a significantly increased risk of death.In china, data indicate [4] that approximately 20.6% of hospitalized patients in critical care medicine departments suffer from sepsis, with mortality rates for sepsis and septic shock reaching 35.5% and 53.3%, respectively. This condition places a substantial strain on medical resources and represents a major threat to public health. Investigating the pathophysiological mechanisms of sepsis and exploring targeted therapeutic strategies are of significant research importance.

Traditional Chinese medicine (TCM) offers significant advantages in the treatment of sepsis and has been increasingly applied in clinical practice in recent years. Studies have confirmed a relationship between the regulation of the NLRP3 inflammasome and the therapeutic effects of TCM in sepsis management [5]. palmatine, an active compound extracted from Phellodendron chinense, Phellodendron chinense bark, and Rhizoma Coptidis [6], has demonstrated promising pharmacological properties.

Modern pharmacological research [7-8] shows that palmatine exhibits antibacterial effects against various pathogens, including Staphylococcus aureus, Escherichia coli, and Streptococcus pneumoniae, as well as antifungal and antiparasitic activities. Additionally, palmatine can reduce inflammatory responses by inhibiting the production of pro-inflammatory cytokines, such as interleukins and tumor necrosis factor, thereby mitigating inflammation. Its antioxidant properties, including free radical scavenging and inhibition of lipid peroxidation, protect cells from oxidative stress-induced damage. In sepsis treatment, palmatine has been found to neutralize endotoxins, reduce the release of interleukin-6 and tumor necrosis factor-α induced by endotoxins, and protect the functions of vital organs [9]. Palmatine, another bioactive compound, has demonstrated efficacy in treating various diseases. Recent studies have highlighted its broad health benefits, including its therapeutic potential in gastric ulcers, skin cancer, and ulcerative colitis. These effects are attributed to its anti-inflammatory and antioxidant properties mediated by the NF-κB/Nrf2 signaling pathway and the regulation of the NLRP3 inflammasome [10-12].

Pyroptosis has recently been recognized as a novel form of programmed cell death, characterized by the release of numerous proinflammatory cytokines, such as IL-1β and IL-18 [13]. The signal transduction mechanisms of pyroptosis include the caspase-1-dependent classical pathway, the caspase-4/5/11-dependent non-classical pathway, and the caspase-3-dependent pathway [14]. All of these pathways share a common feature: the formation of pores in the cell membrane and the release of cellular contents mediated by gasdermins (GSDMs), ultimately leading to cell swelling and rupture. This process triggers a cascade of inflammatory responses [15,16]. Pyroptosis plays a dual role in the immune system. On one hand, it promotes the release of pathogens by destroying infected cells, allowing them to be phagocytosed and cleared by immune cells. This mechanism reduces the antigen burden and helps eliminate intracellular pathogens [17]. On the other hand, in the context of sepsis, moderate pyroptosis serves as a protective mechanism against infection by pathogenic microorganisms. However, excessive activation of pyroptosis can exacerbate sepsis and septic shock [18,19]. Moreover, overactivation of pyroptosis can cause significant organ damage, contributing to the progression of sepsis. Therefore, inhibiting pyroptosis represents a promising therapeutic strategy for treating sepsis [20]. Previous studies have demonstrated that pyroptosis plays a significant role in the progression of sepsis and sepsis-related organ dysfunction. palmatine has been shown to play an important role in the treatment of sepsis. However, whether palmatine exerts its effects on sepsis by modulating pyroptosis remains largely unexplored. This study aims to investigate whether palmatine can influence sepsis by modulating pyroptosis mechanisms through related signaling pathways.

## Materials and methods

### Obtainment and preprocessing of sepsis datasets

The gene expression data of sepsis patients (GSE123729) were obtained from the GEO database (https://www.ncbi.nlm.nih.gov/geo/). Gene probes were converted into gene symbols using the corresponding annotation files provided in the dataset.

### Identification of differentially expressed pyroptosis-related genes

We performed differential analysis on the GSE123729 dataset of sepsis patients and used the “limma” R package in Bioconductor to identify differentially expressed genes in GSE123729 with P < 0.05 and |logFC|> 1. We identified a total of 60 pyroptosis-related genes through the Gene Set Enrichment Analysis (GSEA) website (http://www.gsea-msigdb.org/gsea/index.jsp) and previous literature [21]. By taking the intersection of differentially expressed genes in GSE123729 and pyroptosis-related genes, we obtained pyroptosis-related differentially expressed genes. At the same time, the expression differences of these pyroptosis-related differentially expressed genes were analyzed in the dataset and verified using the GEO dataset.

### Correlation analysis and construction of PPI network

We performed correlation analysis on the differentially expressed genes related to pyroptosis using the rcorr function in the “hmisc” R package and visualized them using the “corrplot” R package. We also constructed a PPI network for the differentially expressed genes related to pyroptosis.

### GO and KEGG enrichment

We performed GO and KEGG enrichment analysis on the differentially expressed genes related to pyroptosis. The biological functions of the identified differentially expressed genes related to pyroptosis were evaluated using Gene Ontology (GO) functional enrichment analysis and Kyoto Encyclopedia of Genes and Genomes (KEGG) pathway analysis and the “clusterProfiler (4.10.1)” R package.

### Consensus clustering

Consensus clustering, a method for identifying molecular subtypes based on the approximate number of clusters, was used to discover sepsis subgroups associated with the expression of pyroptosis-related regulators by the k-means method. The maximum number of categories to be evaluated was 10, and the number of iterations for each k was 100. The euclidean distance was chosen as the cluster distance. The number of clusters was calculated by the consensus clustering algorithm using the “ConsensuClusterPlus (1.66.0)” R package. The graphical results are heatmaps of consensus clustering.

### Immune infiltration analysis

We used the CIBERSORT algorithm to calculate the immune infiltration scores of sepsis patients between subtypes after consensus clustering to confirm the relevant proportions of immune infiltrating cells.

### Gene set variation analysis (GSVA)

GSVA analysis is one of the GSEA algorithms that can explore the differences in biological pathways between different pattern clusters based on enrichment scores. The “GSVA (1.16.0)” R package was applied to perform functional enrichment analysis on sepsis disease samples in GSE123729 to obtain enriched pathways. We downloaded “c2.cp.kegg.v7.4.symbols.gmt” from the MsigDB database for analysis. Corrected P values < 0.05 were considered to indicate statistical differences between different clusters.

### GO and KEGG enrichment between clusters

We performed differential expression analysis on genes between two different clusters in GSVA enrichment analysis, with the screening conditions of P < 0.05, log2 (Fold Change) > 0.5. Then GO and KEGG enrichment analysis were performed on the screened differential genes. The biological functions of differentially expressed genes identified in different clusters were evaluated using Gene Ontology (GO) functional enrichment analysis and Kyoto Encyclopedia of Genes and Genomes (KEGG) pathway analysis with the “clusterProfiler (4.10.1)” R package. For GO analysis, enriched biological processes (BPs), molecular functions (MFs), and cellular components (CCs) were evaluated.

### Network pharmacology and palmatine-related pathways

We used the Traditional Chinese Medicine Systems Pharmacology Database and Analysis Platform (TCMSP: http://tcmspw.com/tcmsp.php) to identify TCMs containing palmatine as active ingredients and their related targets. The screening criteria were established based on two basic pharmacokinetic indicators: oral bioavailability (OB) and drug similarity (DL), with thresholds set at OB ≥ 30% and DL ≥ 0.18. We integrated the targets corresponding to the compound name palmatine in different TCMs to form a complete target list, and then converted the targets into gene symbols. After that, the gene symbols can be converted into IDs, GO, KEGG enrichment analysis can be performed, and the relationship between genes and KEGG pathways can be visualized to construct a gene-KEGG pathway network.

### Qpcr experiments to verify gene expression

Total RNA was extracted using a Total RNA Extractor (Hangzhou Haoke Biotechnology Co., Ltd., HKR022). The extracted RNA was reverse transcribed into cDNA using the All-in-One First-Strand cDNA Synthesis SuperMix (TRAN, AE341), and real-time fluorescent quantitative PCR was performed using 2X SYBR Green qPCR Master Mix (APExBIO, K1070). The expression fold changes were calculated using the ΔΔCT method, with an internal reference gene as the control.

## Results

### Expression and correlation analysis of pyroptosis-related genes in sepsis

We identified 60 pyroptosis-related genes using the Gene Set Enrichment Analysis (GSEA) website (http://www.gsea-msigdb.org/gsea/index.jsp) and previous literature [17]. Using the GEO dataset GSE123729, we analyzed the differential expression of these 60 pyroptosis-related genes in sepsis patients compared to controls, ultimately identifying 9 differentially expressed genes: CXCL8, PTGS2, NLRP1, IL1B, AKT1, IRF1, CASP5, NLRP6, and NLRC4 (Fig 1B). A heat map displaying the expression levels of these differentially expressed genes in GSE123729 is shown in Fig 1A.

**Fig 1A.**
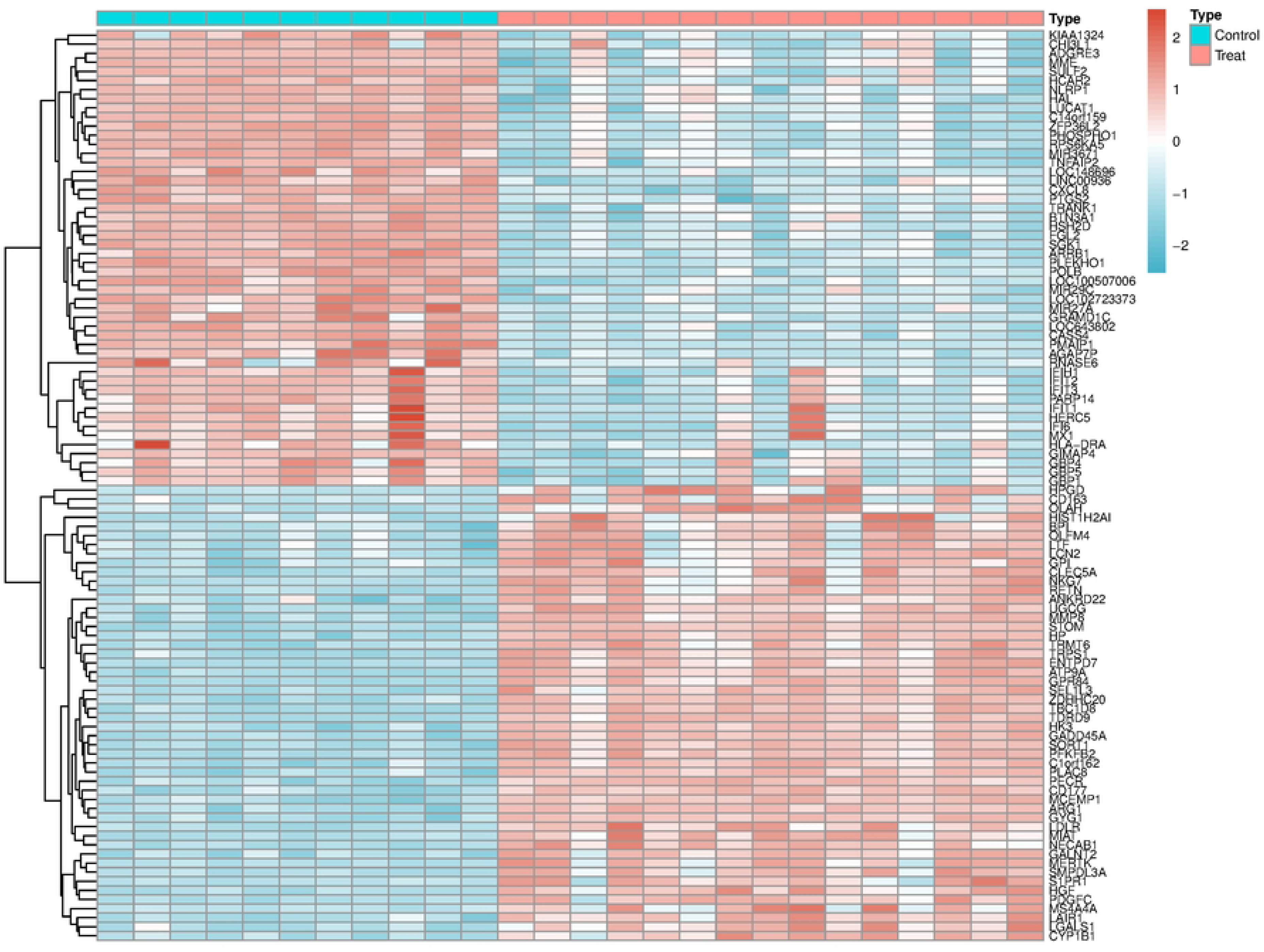
Heatmap of DEGs between sepsis and control groups.

**Fig 1B.**
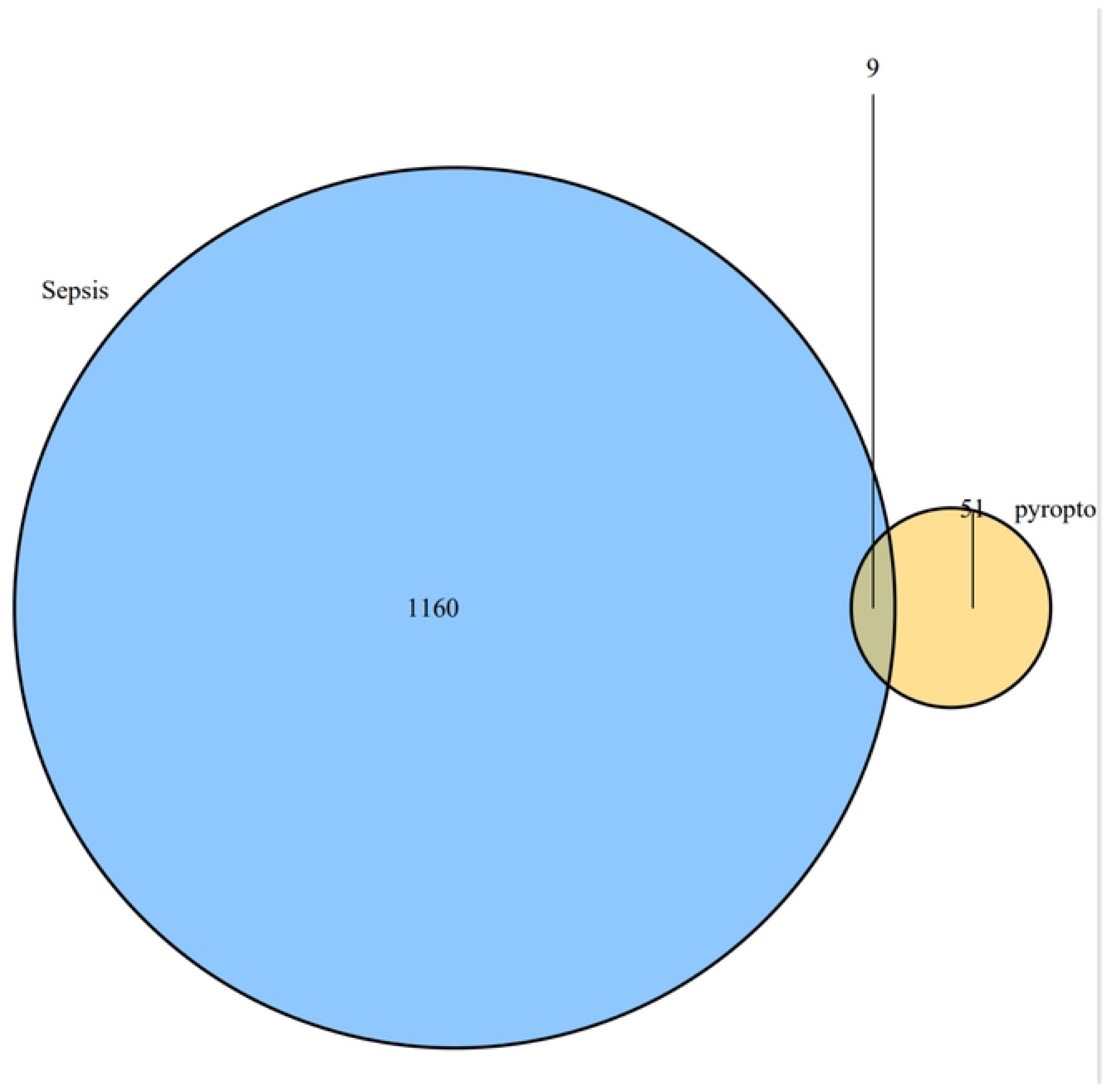
Intersection of differentially expressed genes and pyroptosis-related genes.

To further explore the interactions among these pyroptosis-related genes, we performed a protein-protein interaction (PPI) analysis using the STRING platform, with the interaction score threshold set at 0.400 (Fig 1C). Additionally, we conducted a correlation analysis (Fig 1D), revealing that NLRP family genes (NLRP1, NLRP4, NLRP6) were closely interconnected, particularly with CASP5 and IL1B, while CXCL8 exhibited weaker associations with IRF1 and minimal interactions with the NLRP family genes.

**Fig 1C.**
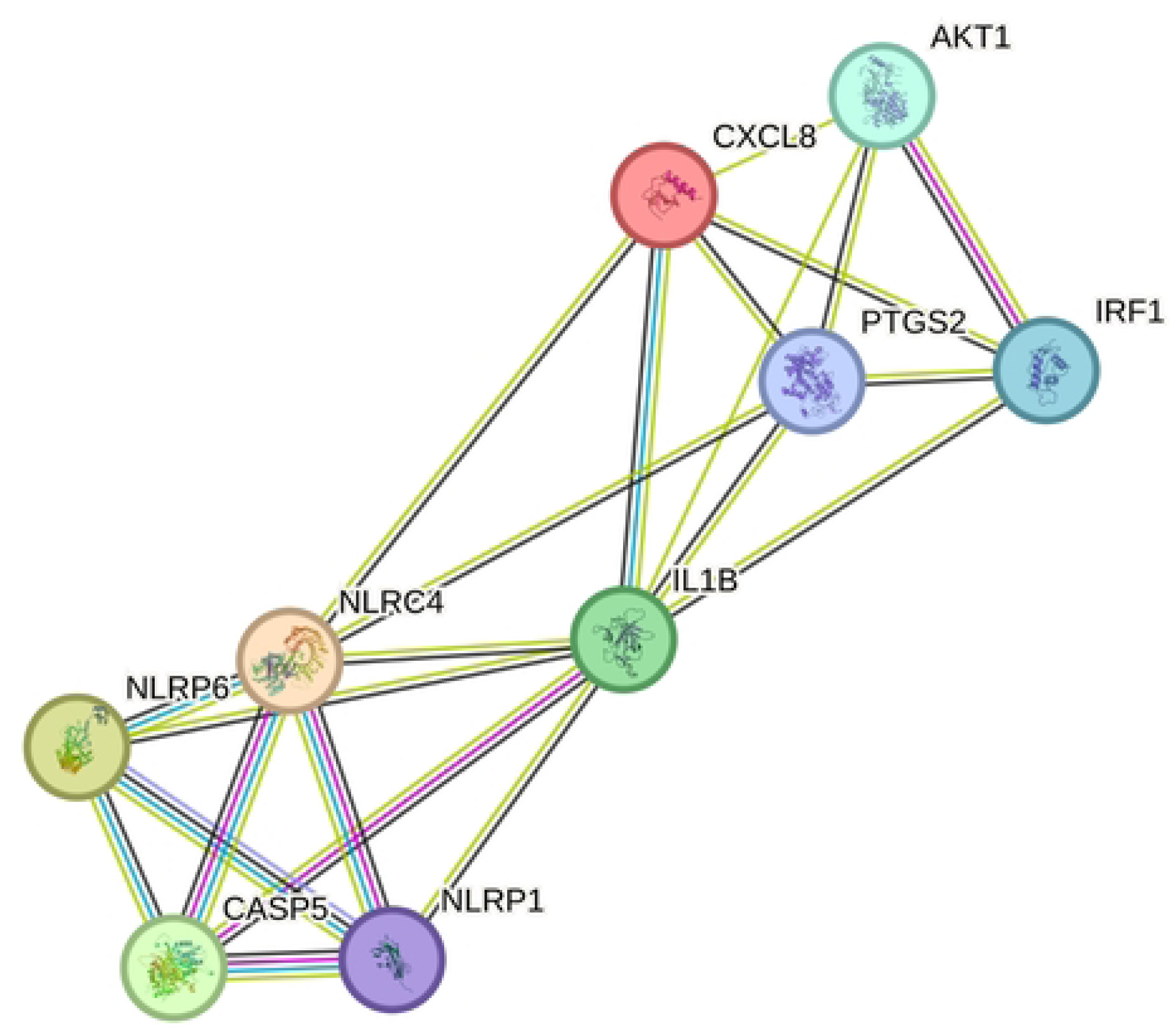
PPI netwoek of 9 intersection genes.

**Fig 1D.**
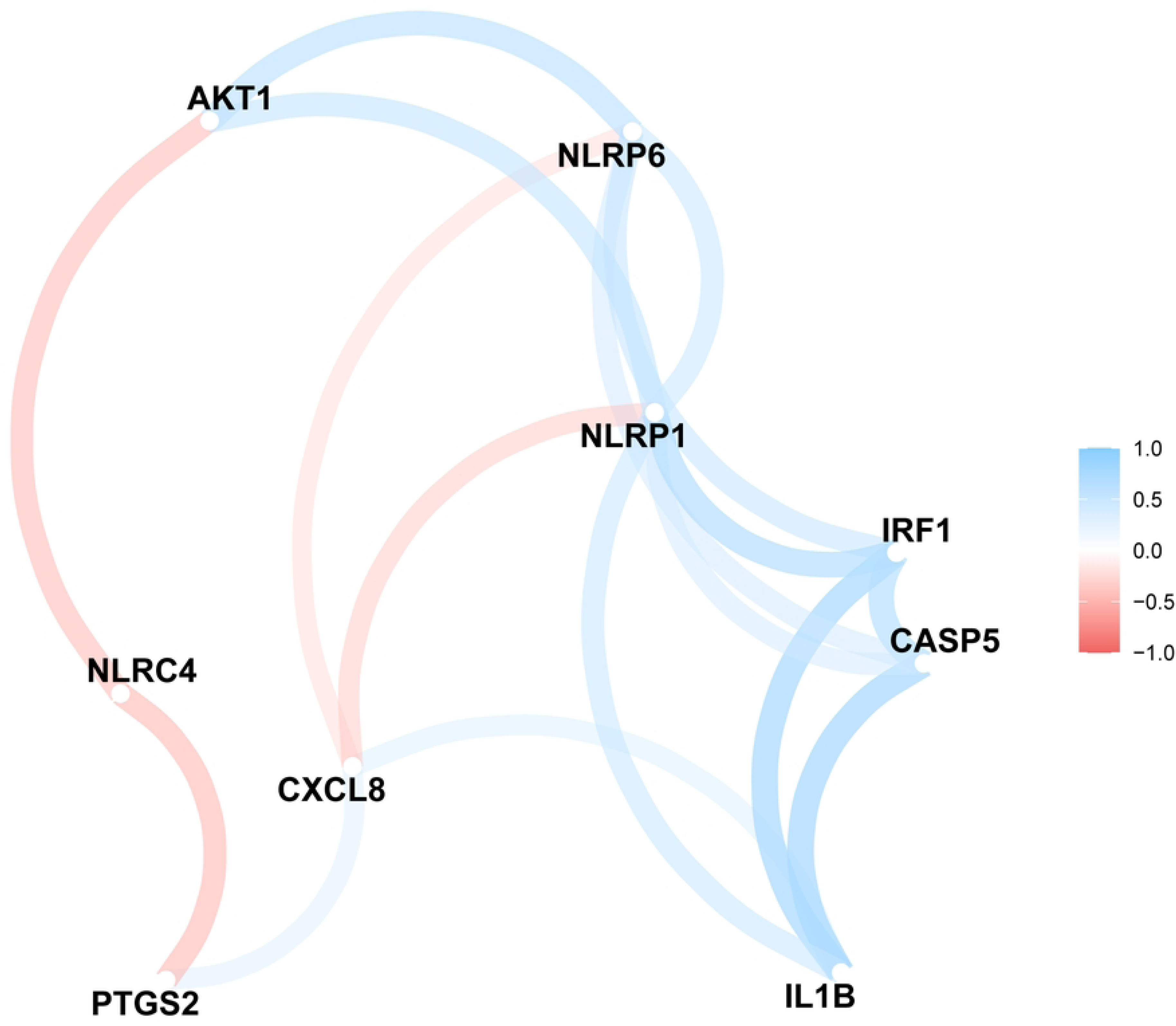
Correlation of 9 intersection genes.

We performed Gene Ontology (GO) and KEGG enrichment analyses on these 9 differentially expressed pyroptosis-related genes (Figs 1E, 1F). The GO enrichment analysis revealed significant enrichment in: Biological Processes (BP): Positive regulation of nitric oxide metabolic process, Positive regulation of nitric oxide biosynthetic process, Pyroptosis, and Regulation of inflammatory response. Cellular Component (CC): Canonical inflammasome complex. Molecular Function (MF): Cysteine-type endopeptidase activity involved in apoptotic process, Cytokine activity, and Pattern recognition receptor activity. To visualize the expression differences, we constructed a box plot of these 9 pyroptosis-related genes in sepsis patients and controls (Fig 1G). The analysis showed that NLRC4 was highly expressed and upregulated in sepsis patients compared to controls, whereas the remaining 8 genes (CXCL8, PTGS2, NLRP1, IL1B, AKT1, IRF1, CASP5, NLRP6) were downregulated in sepsis patients. To validate our findings, we analyzed three additional sepsis-related GEO datasets: GSE57065, GSE95233, and GSE54514 (Figs 1H, 1I, 1J). These validation sets confirmed the observed gene expression patterns, showing that NLRC4 was upregulated while the remaining genes were consistently downregulated in sepsis patients.

**Fig 1E.**
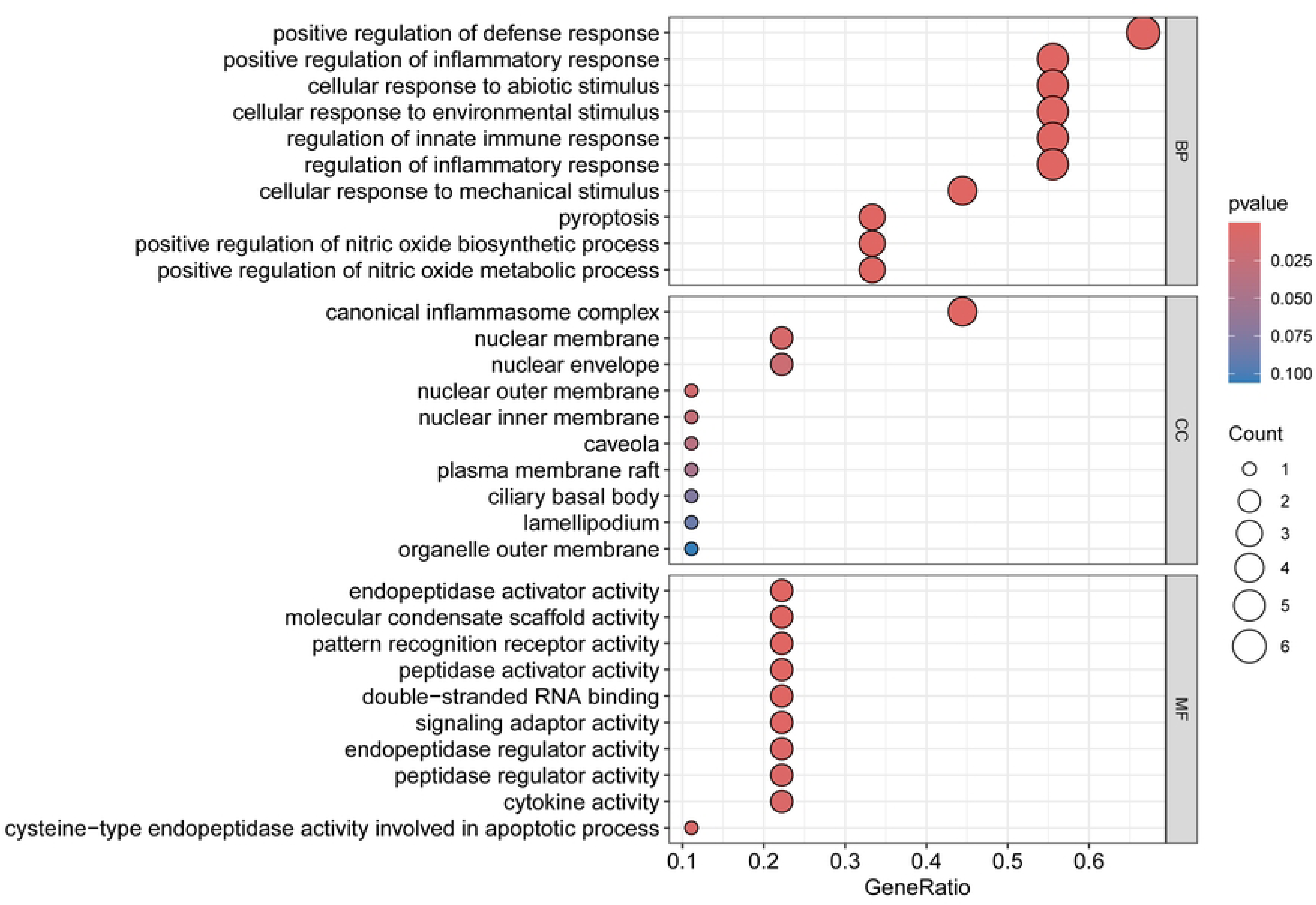
GO enrichment of 9 intersection genes.

**Fig 1F.**
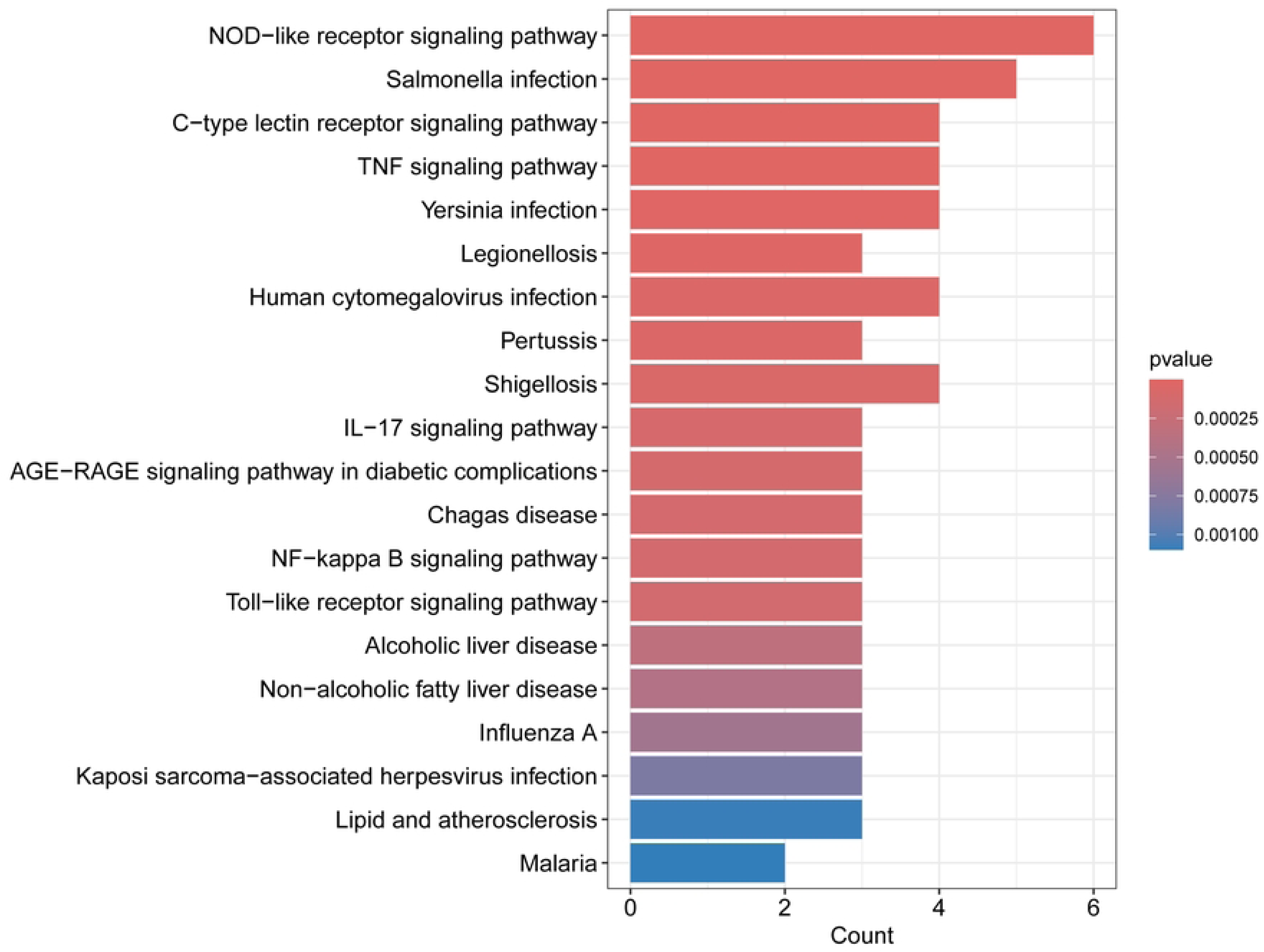
KEGG enrichment of 9 intersection genes.

**Fig 1G.**
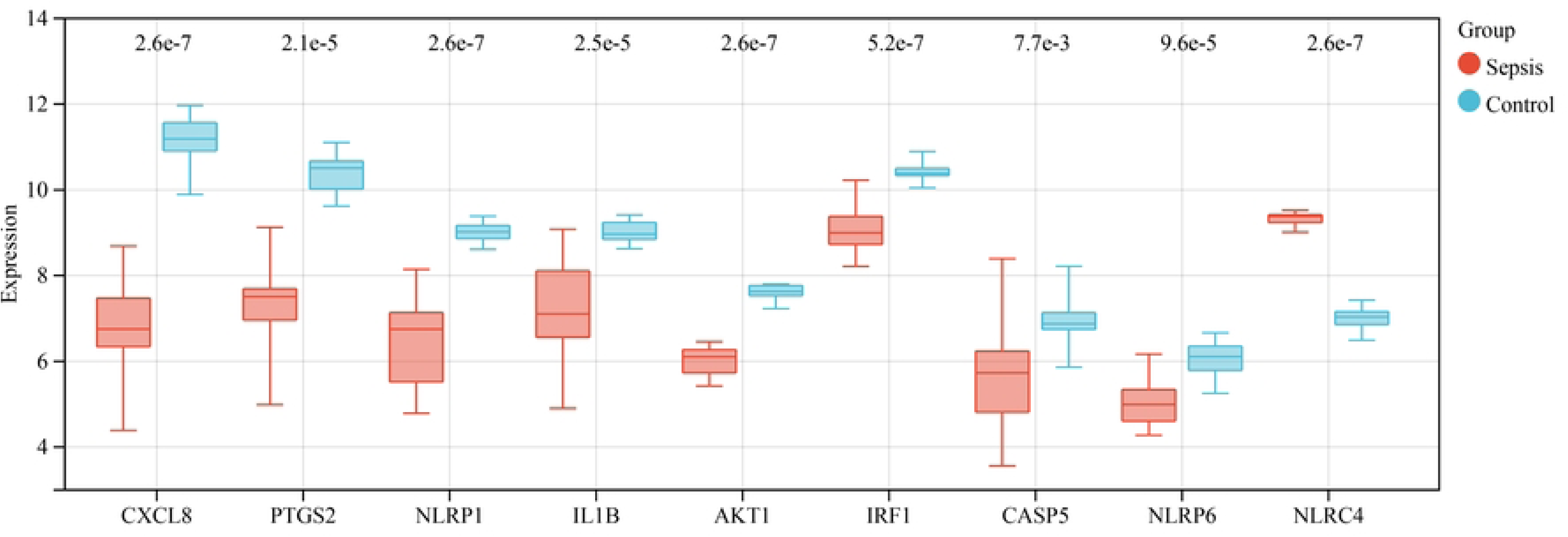
Expression level of 9 intersection genes.

**Fig 1H.**
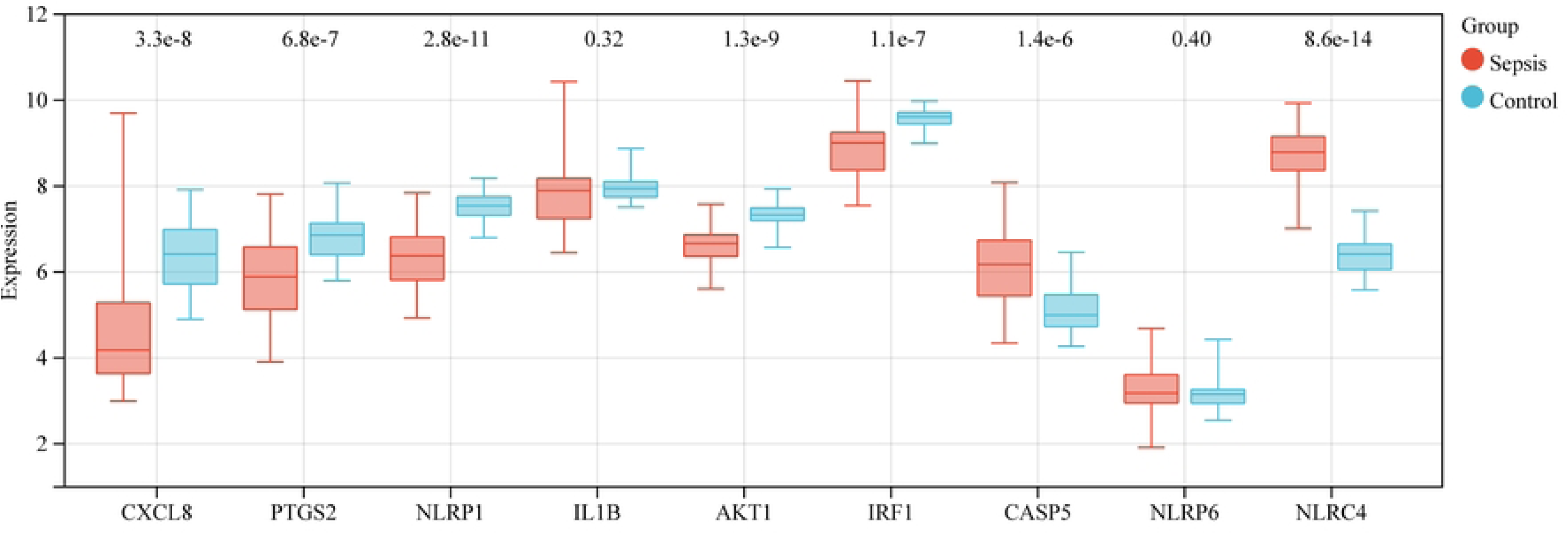
Expression level of 9 intersection genes in GSE57065.

**Fig 1I.**
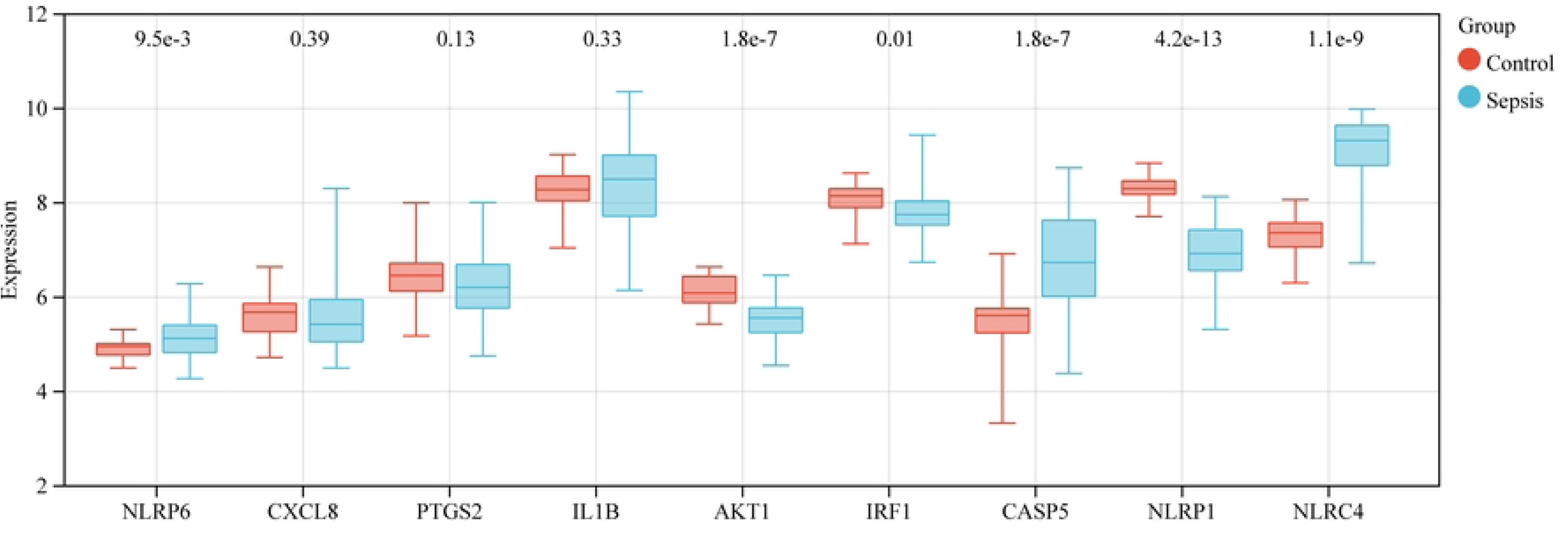
Expression level of 9 intersection genes in GSE95233.

**Fig 1J.**
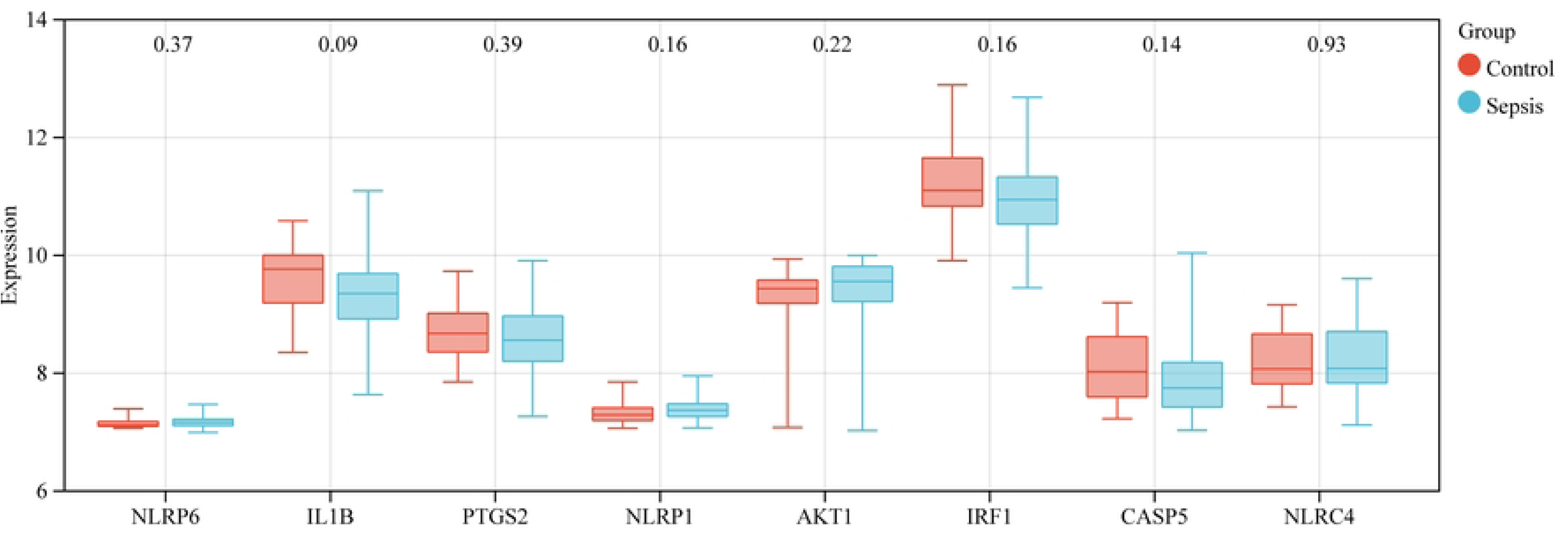
Expression level of 9 intersection genes in GSE54514.

### Differences in pyroptosis-related molecular subsets and immune microenvironment between subgroups

To investigate the relationship between the 9 pyroptosis-related differentially expressed genes and sepsis phenotypes, we conducted a consistent clustering analysis on 15 sepsis patients from the GSE123729 dataset. When the cluster variable k was set to 2, the intra-group correlation was maximized, and the inter-group correlation was minimized (Figs 2A, 2B). This result indicates that the 15 patients can be effectively divided into two distinct clusters based on the 9 pyroptosis-related differentially expressed genes. Seven patients were classified into Cluster A, while eight patients were grouped into Cluster B, with their inter-group gene expression visualized in a heat map (Fig 2C). To further characterize the immune landscape of the two clusters, we employed the CIBERSORT algorithm to calculate immune infiltration scores for the 15 sepsis patients. The scores revealed notable differences in the proportions of immune infiltrating cells between the two clusters. Specifically, plasma cells exhibited a higher score in Cluster A, while CD4 naïve T cells had a higher score in Cluster B (Fig 2D).

**Fig 2A.**
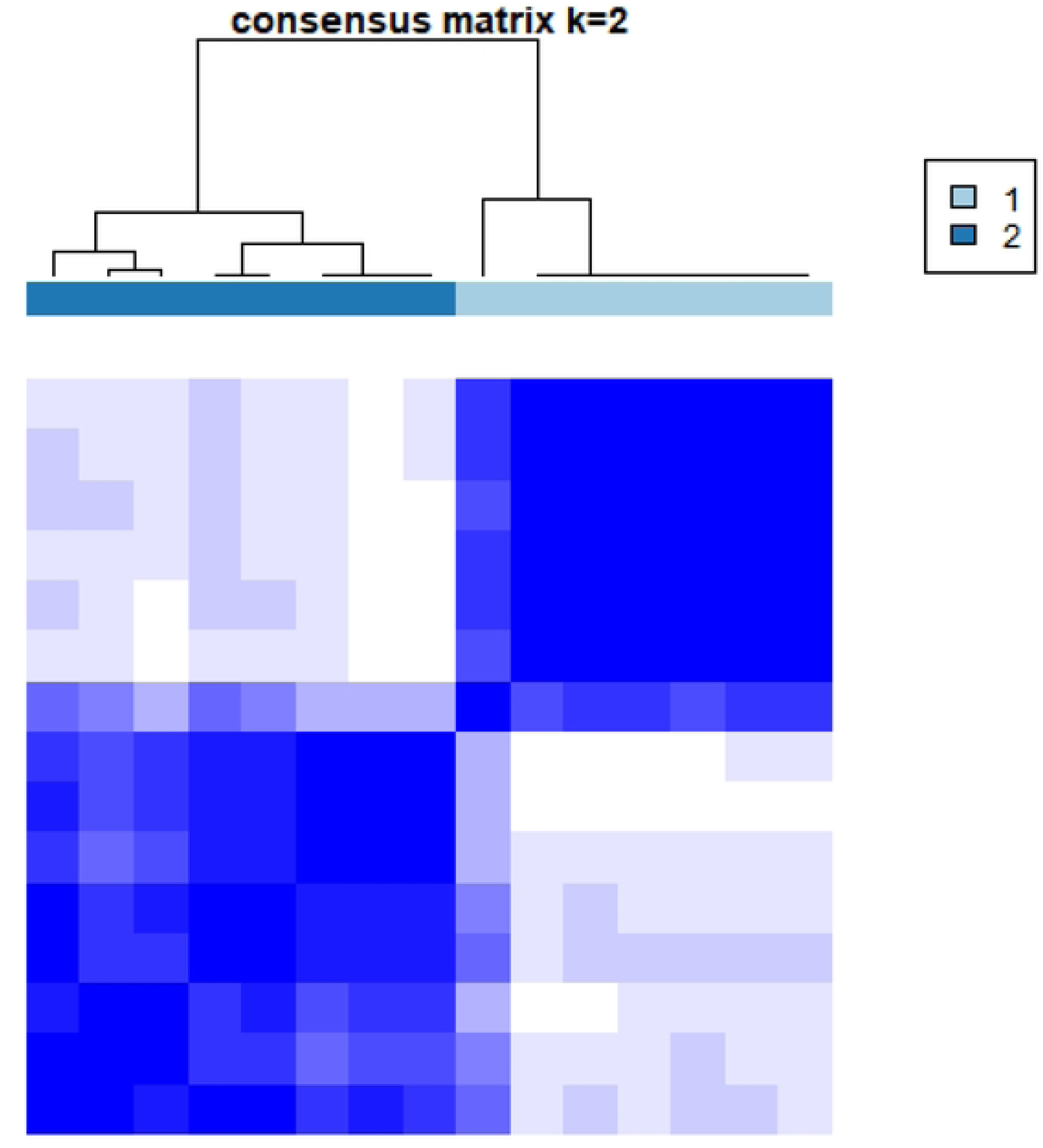
Consensus matrix when k=2.

**Fig 2B.**
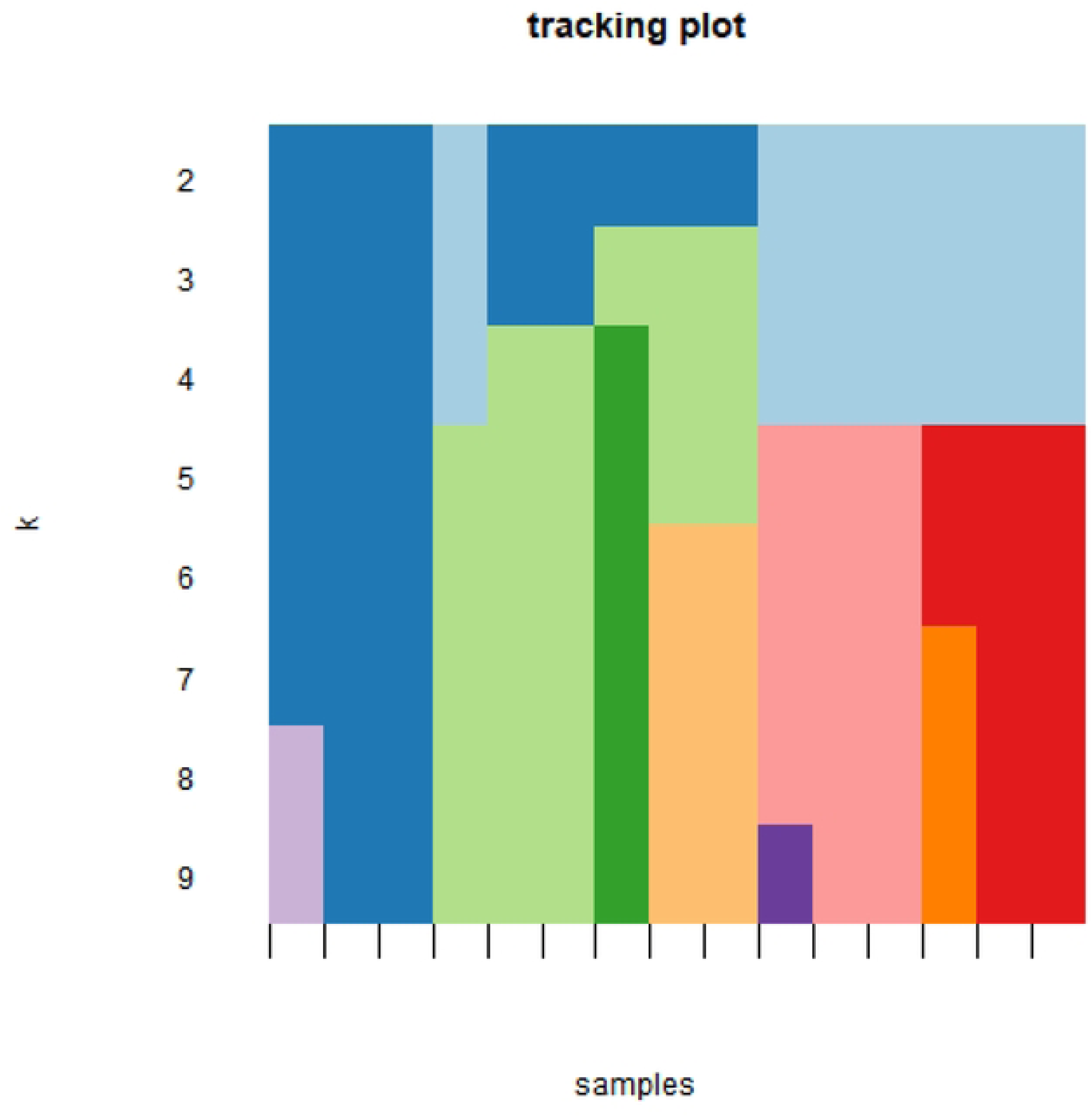
Tracking plot when k=2.

**Fig 2C.**
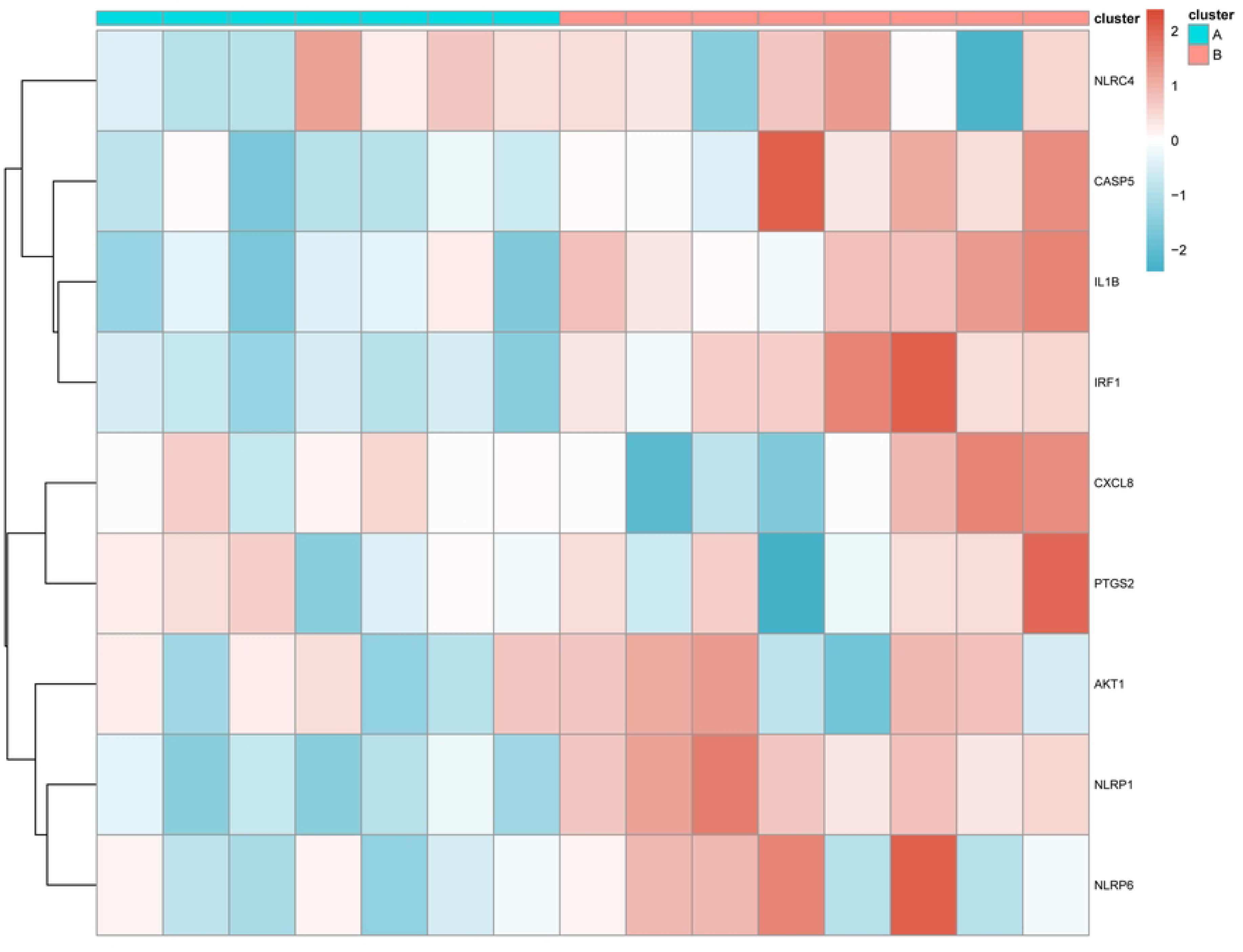
Heatmap of 9 intersection genes of two pyroptosis-related clusters.

**Fig 2D.**
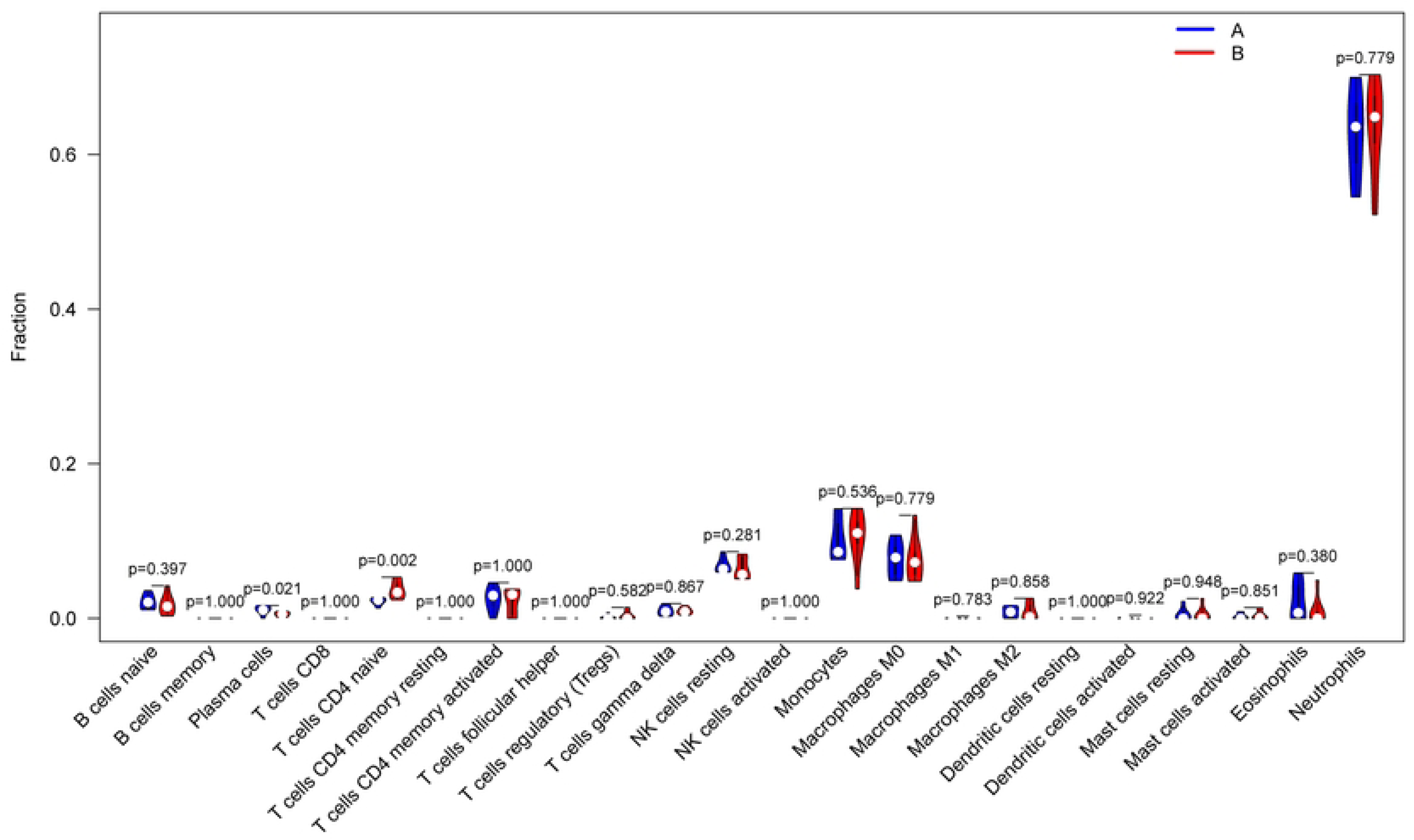
Immune cell infiltration scores of two pyroptosis-related clusters.

### Functional analysis of different molecular subtypes

To investigate the relationship between different model organism clusters mediated by pyroptosis, we performed GSVA enrichment analysis. Our findings revealed that pathways such as KEGG_LYSINE_DEGRADATION and KEGG_MISMATCH_REPAIR were enriched in Cluster A (Cluster A is shown in red, Cluster B in blue), indicating that these pathways are more active in Cluster A. Conversely, pathways such as KEGG_ETHER_LIPID_METABOLISM and KEGG_CALCIUM_SIGNALING_PATHWAY were more enriched in Cluster B (Cluster A is shown in blue, Cluster B in red), indicating higher activity in Cluster B. Overall, Cluster A was primarily enriched in pathways related to immunity and DNA repair, while Cluster B was mainly enriched in pathways associated with signal transduction and metabolism (Fig 3A). Subsequently, we identified differentially expressed genes (DEGs) between the two clusters, which are presented in a heat map (Fig 3B). GO and KEGG pathway analyses were performed on these DEGs. The GO enrichment results were categorized into biological processes (BP), cellular components (CC), and molecular functions (MF), encompassing 21 CC terms and 38 MF terms. The enrichment results are shown in Fig 3C, 3D, 3E, and 3F.

**Fig 3A.**
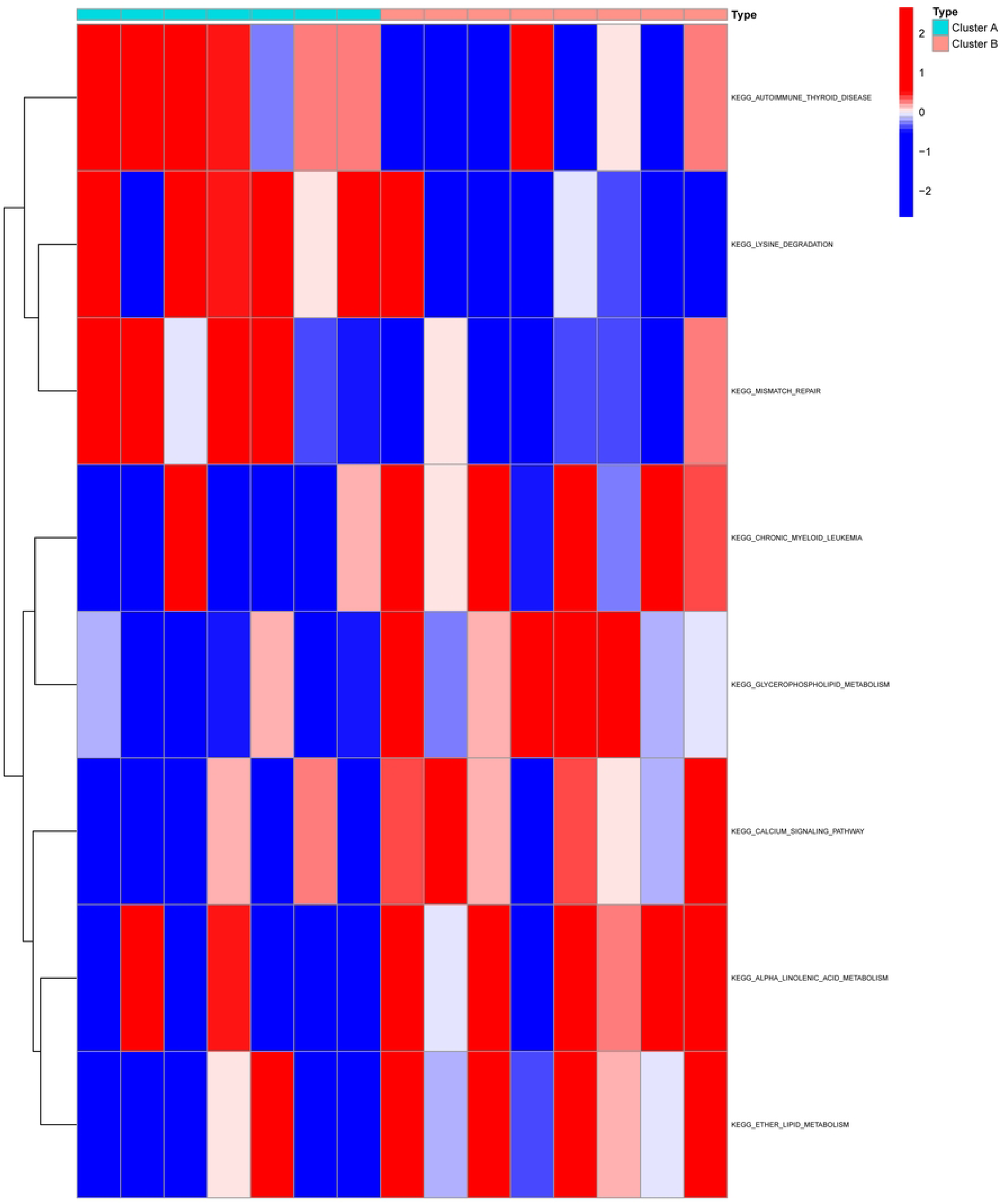
GSVA enrichment of two pyroptosis-related clusters.

**Fig 3B.**
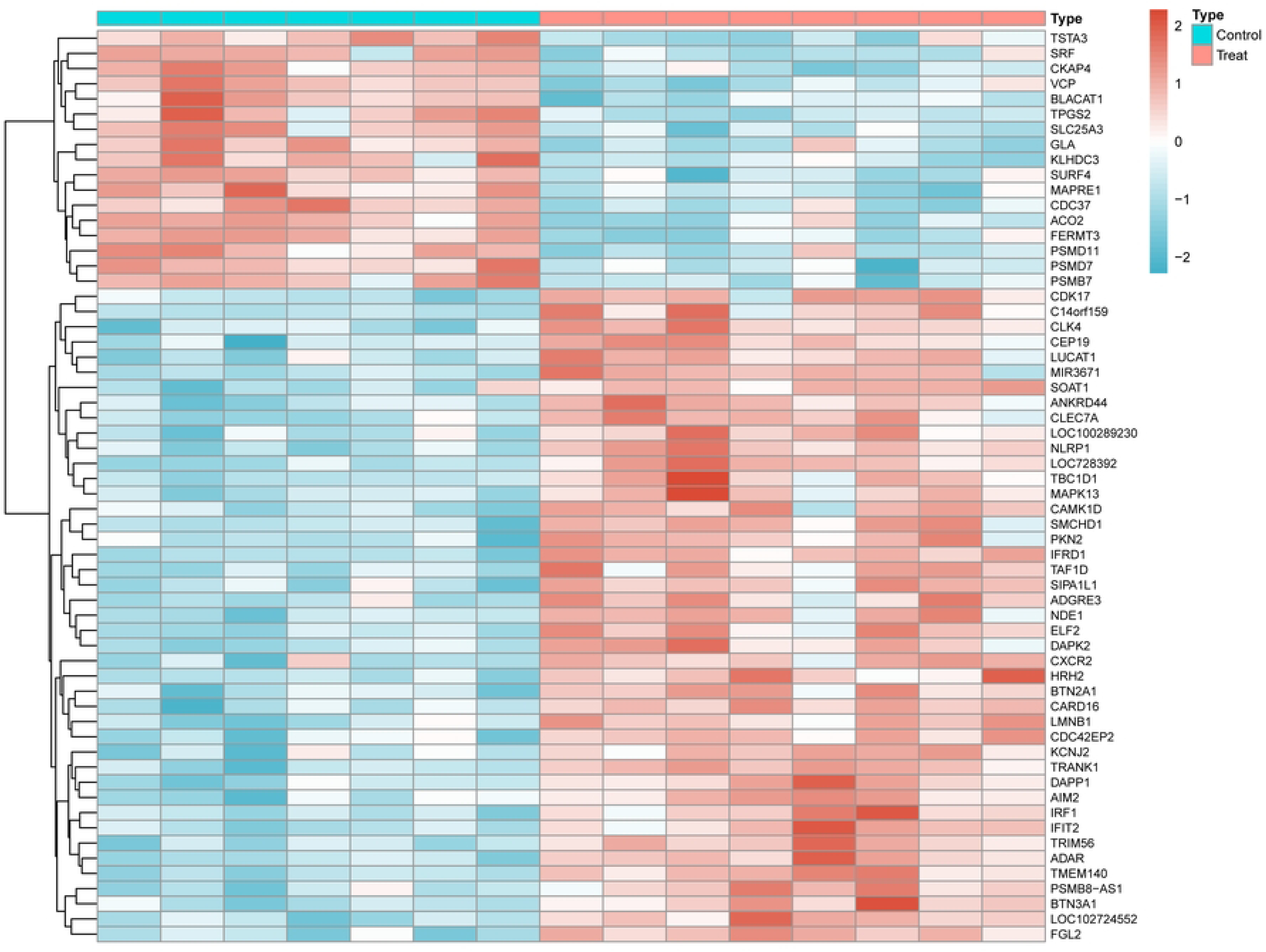
Heamtap of DEGs of two pyroptosis-related clusters.

**Fig 3C.**
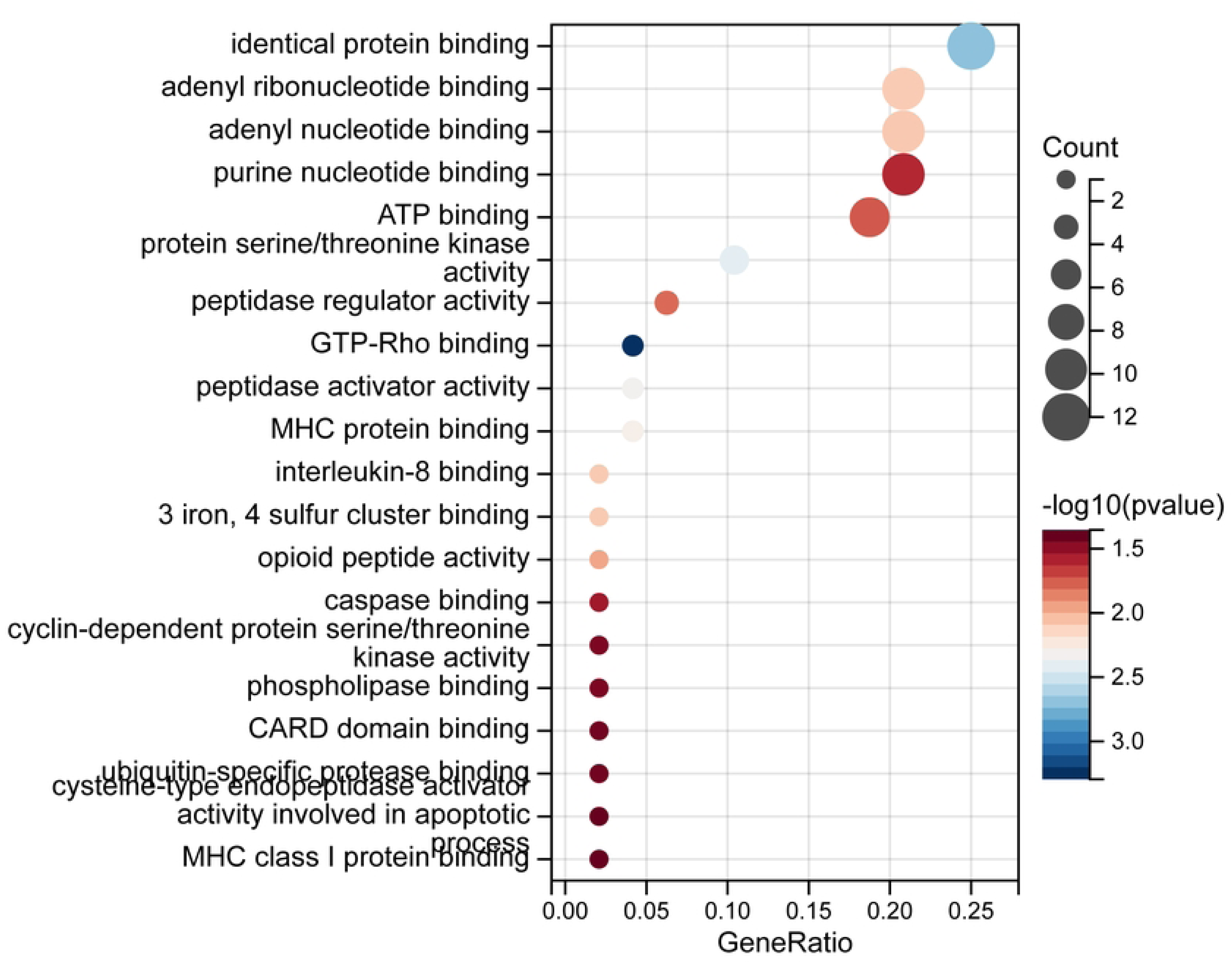
Barplot of BP.

**Fig 3D.**
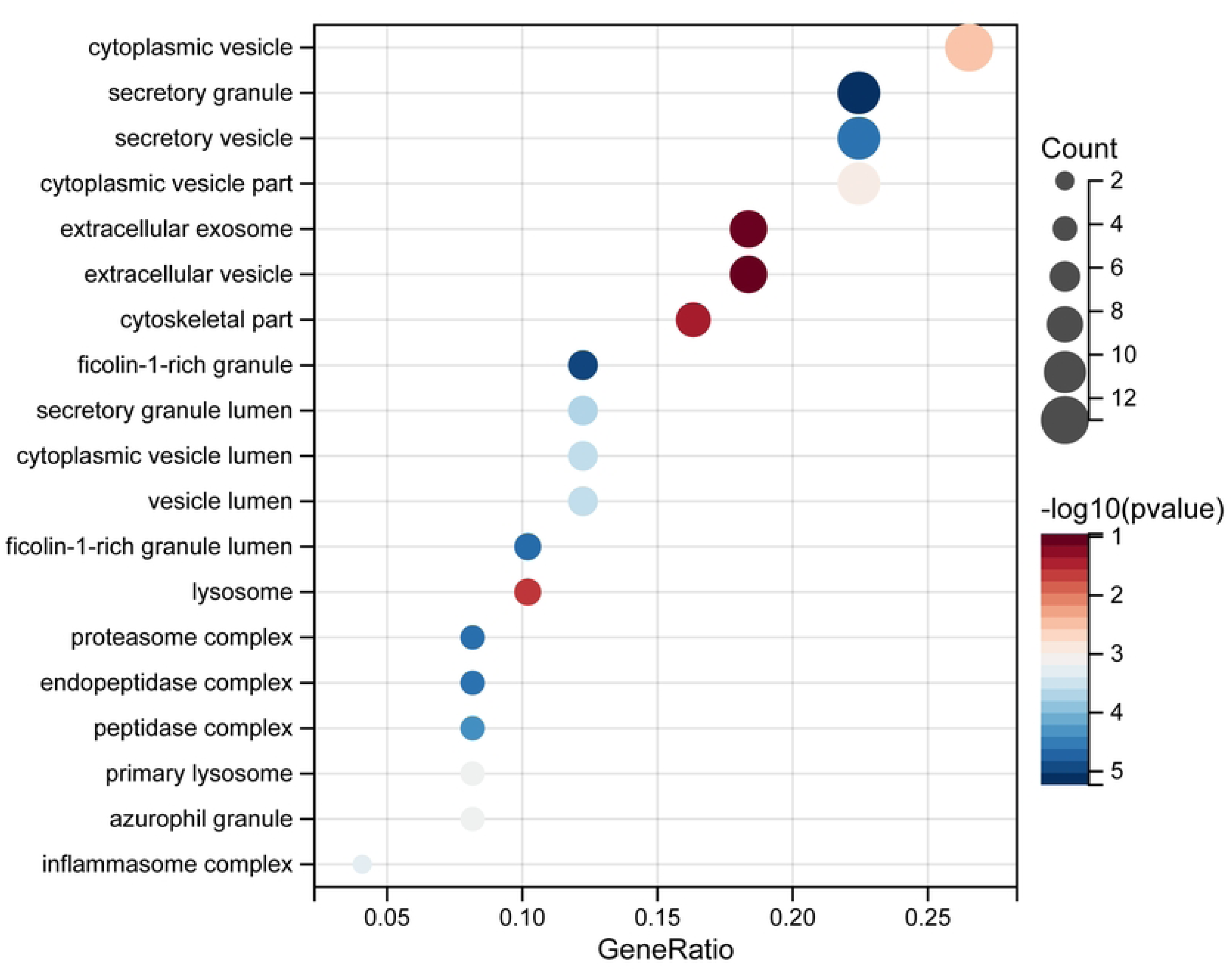
Barplot of CC.

**Fig 3E.**
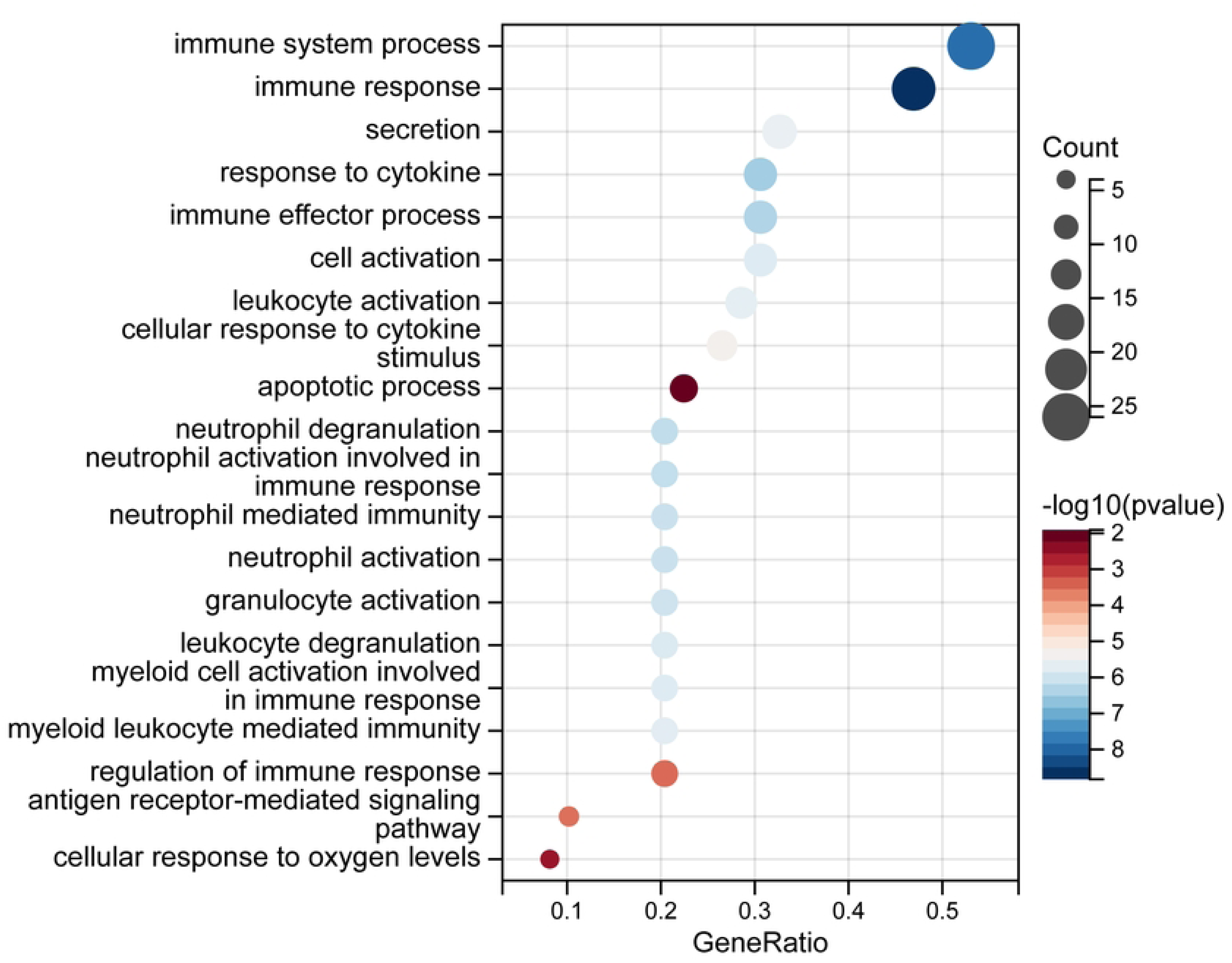
Barplot of MF.

**Fig 3F.**
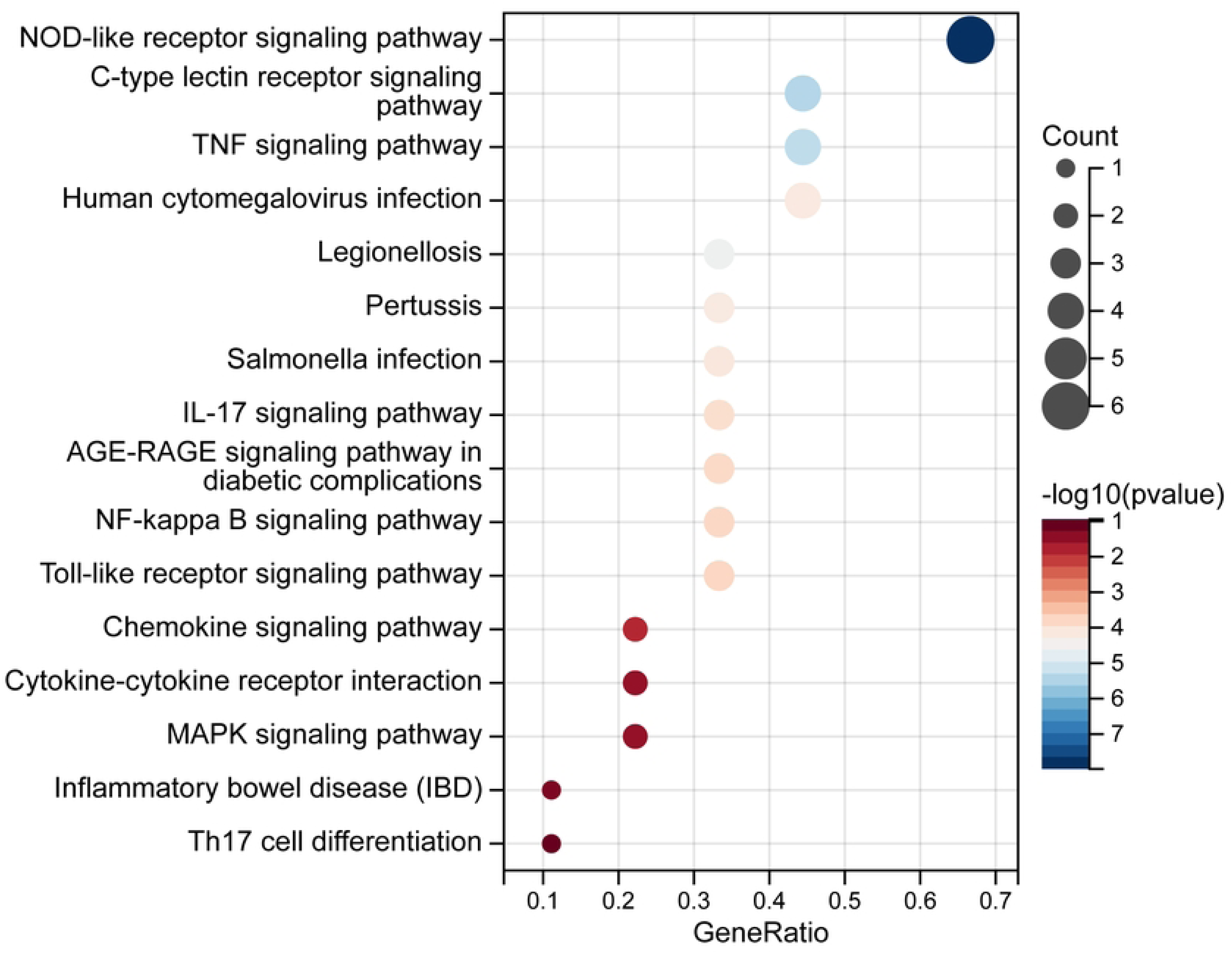
KEGG enrichment of DEGs of two pyroptosis-related clusters.

In the BP category: The most significantly enriched terms included Identical protein binding, Adenyl nucleotide binding, and ATP binding. In the CC category: The most significantly enriched terms were Cytoplasmic vesicle, Cytoplasmic vesicle part, Secretory granule, and Secretory vesicle. In the MF category: Significant terms included Immune system process, Immune response, Response to cytokine, and Cellular response to cytokine stimulus.

For the KEGG signaling pathways, the most significantly enriched pathways included the NOD-like receptor signaling pathway, C-type lectin receptor signaling pathway, TNF signaling pathway, and NF-kappa B signaling pathway.

### Identification of palmatine targets and related pathways by network pharmacology

We identified the relevant targets corresponding to palmatine, a compound in traditional Chinese medicine, and performed GO and KEGG enrichment analyses on these palmatine-related targets (Figs 4A, 4B). Through GO enrichment analysis, we found significant enrichment in terms such as Cellular response to estrogen stimulus, Response to estradiol, Caveola, Membrane raft, Transcription coregulator binding, and RNA polymerase II-specific DNA-binding transcription factor binding. KEGG enrichment analysis revealed significant enrichment in pathways such as the IL-17 signaling pathway, Arachidonic acid metabolism, and Calcium signaling pathway. To further explore the relationship between palmatine-related targets and KEGG pathways, we constructed a target-pathway interaction network (Fig 4C). Within this network, PTGS2 was found to be closely associated with pathways such as the TNF signaling pathway, NF-kappa B signaling pathway, Arachidonic acid metabolism, and IL-17 signaling pathway. Similarly, PTGS1 was identified as being closely related to the Arachidonic acid metabolism signaling pathway.

**Fig 4A.**
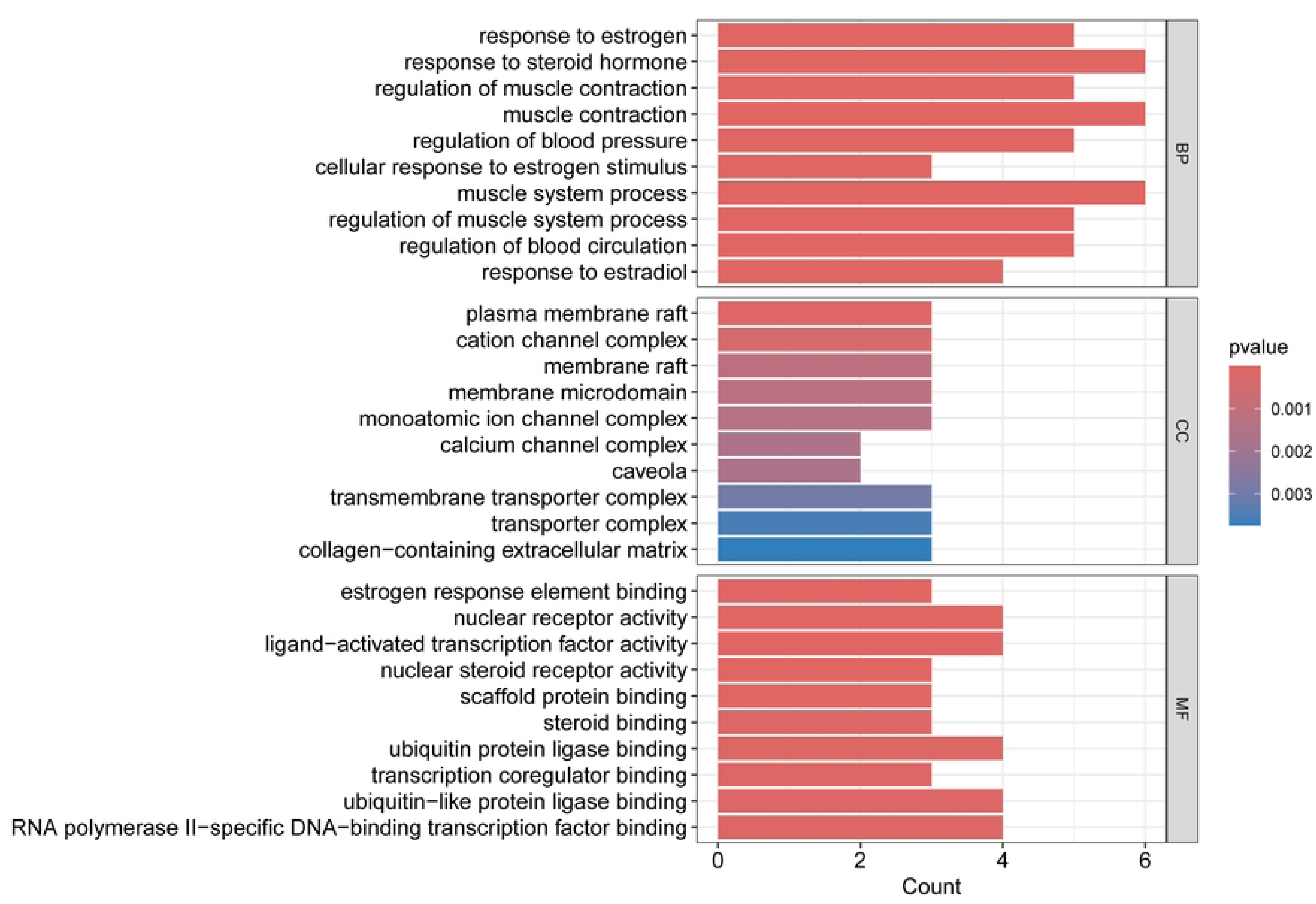
GO enrichment of palmatine targets.

**Fig 4B.**
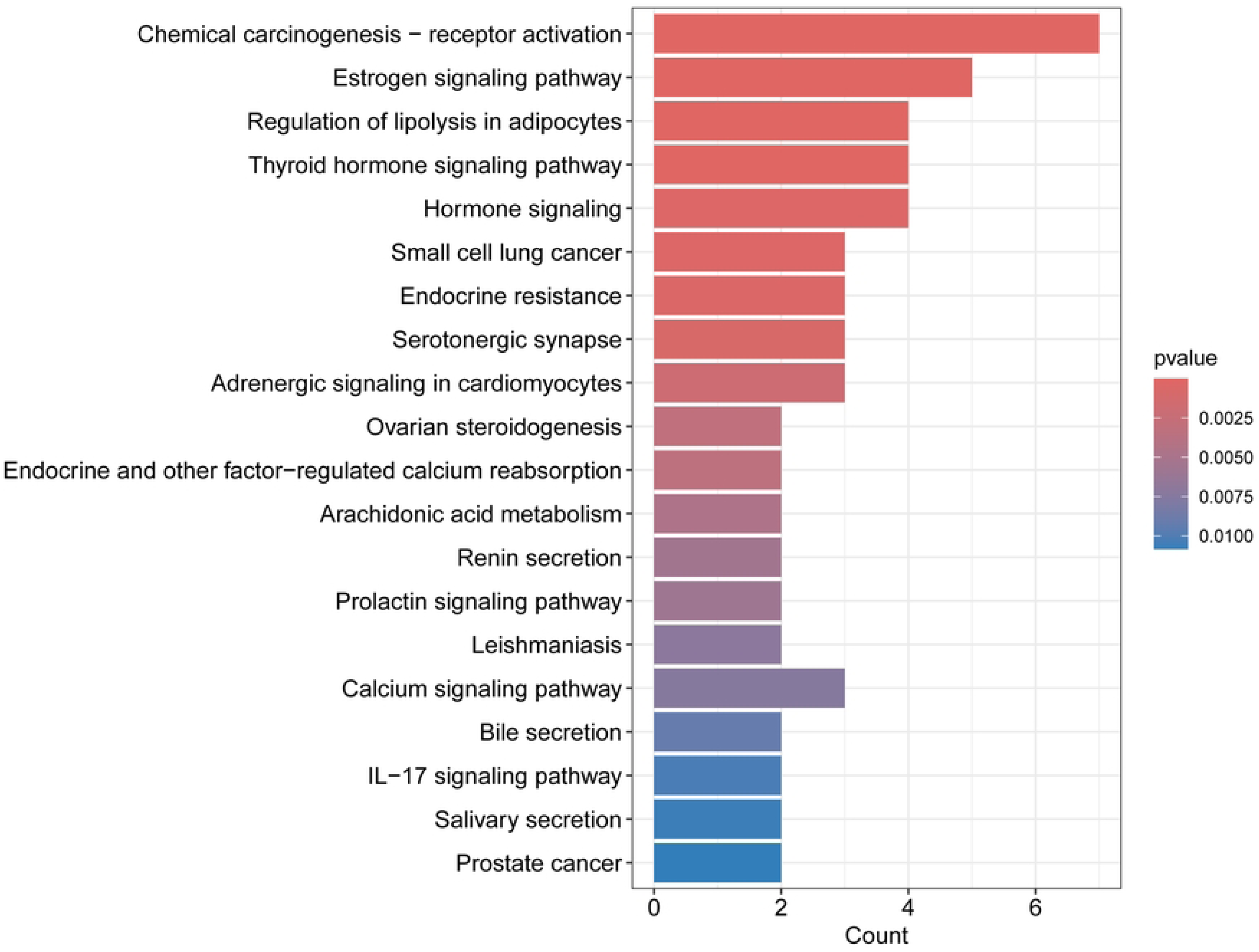
KEGG enrichment of palmatine targets.

**Fig 4C.**
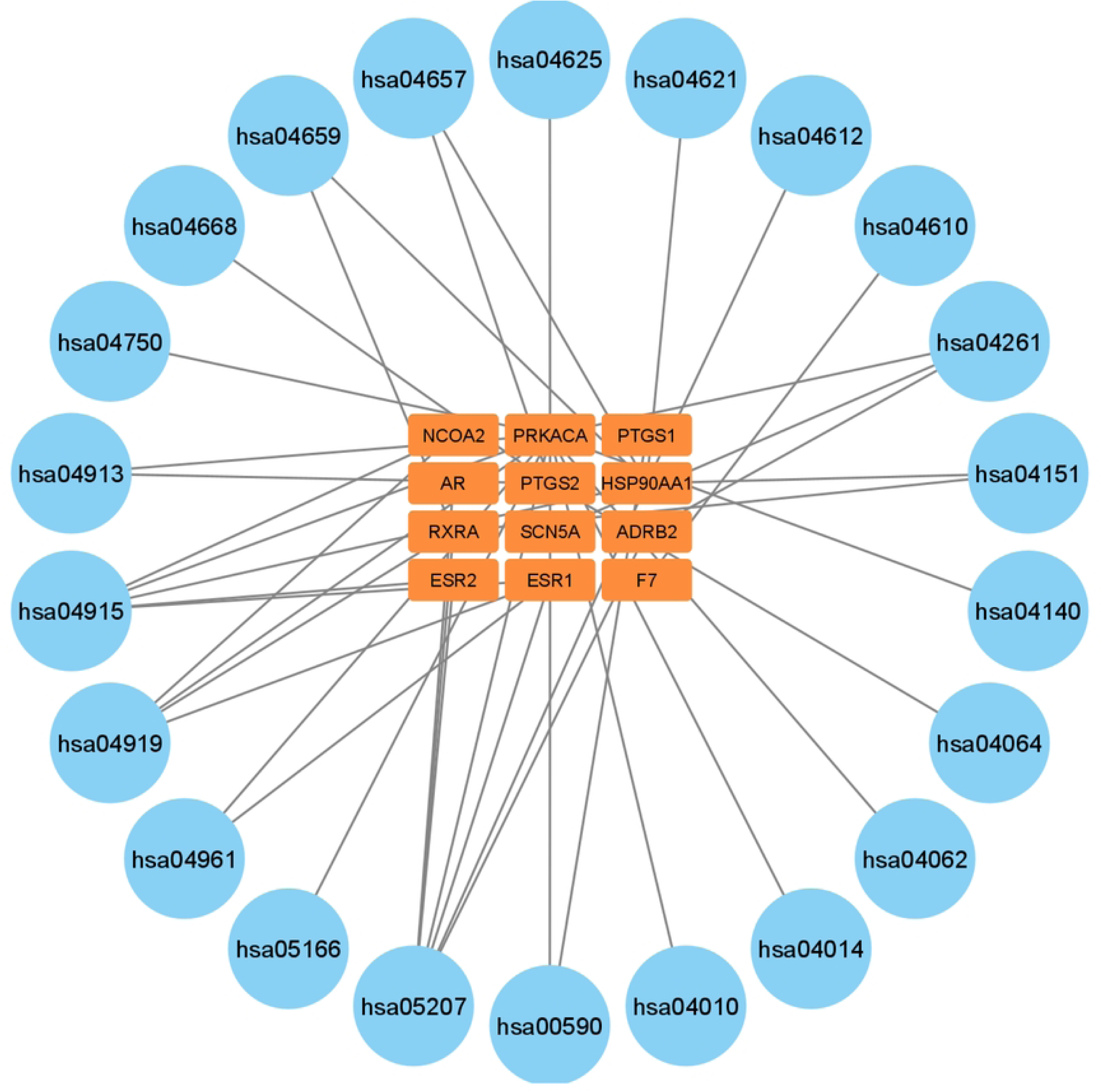
Network of palmatine targets and KEGG pathways.

### Q-PCR Analysis Validation

To further validate the expression of genes in sepsis, we collected serum samples from 5 sepsis and 7 controls for quantitative real-time PCR (qPCR).

We conducted qPCR experiments on selected genes to validate their differential expression between sepsis and control groups. For these experiments, we focused on genes closely associated with the NF-κB signaling pathway, NOD-like receptor signaling pathway, TNF signaling pathway, and PI3K-Akt signaling pathway, which are also relevant targets of bamipine. Specifically, we selected five genes: PRKACA, PTGS2, HSP90AA1, PTPN22 and NLRP3. The qPCR results revealed that these genes were downregulated in patients with sepsis compared to the control group except NLRP3(Figs 5A-5D), and Fig5E NLRP3 was upregulated in patients with sepsis compared to the control group.

**Fig 5A.**
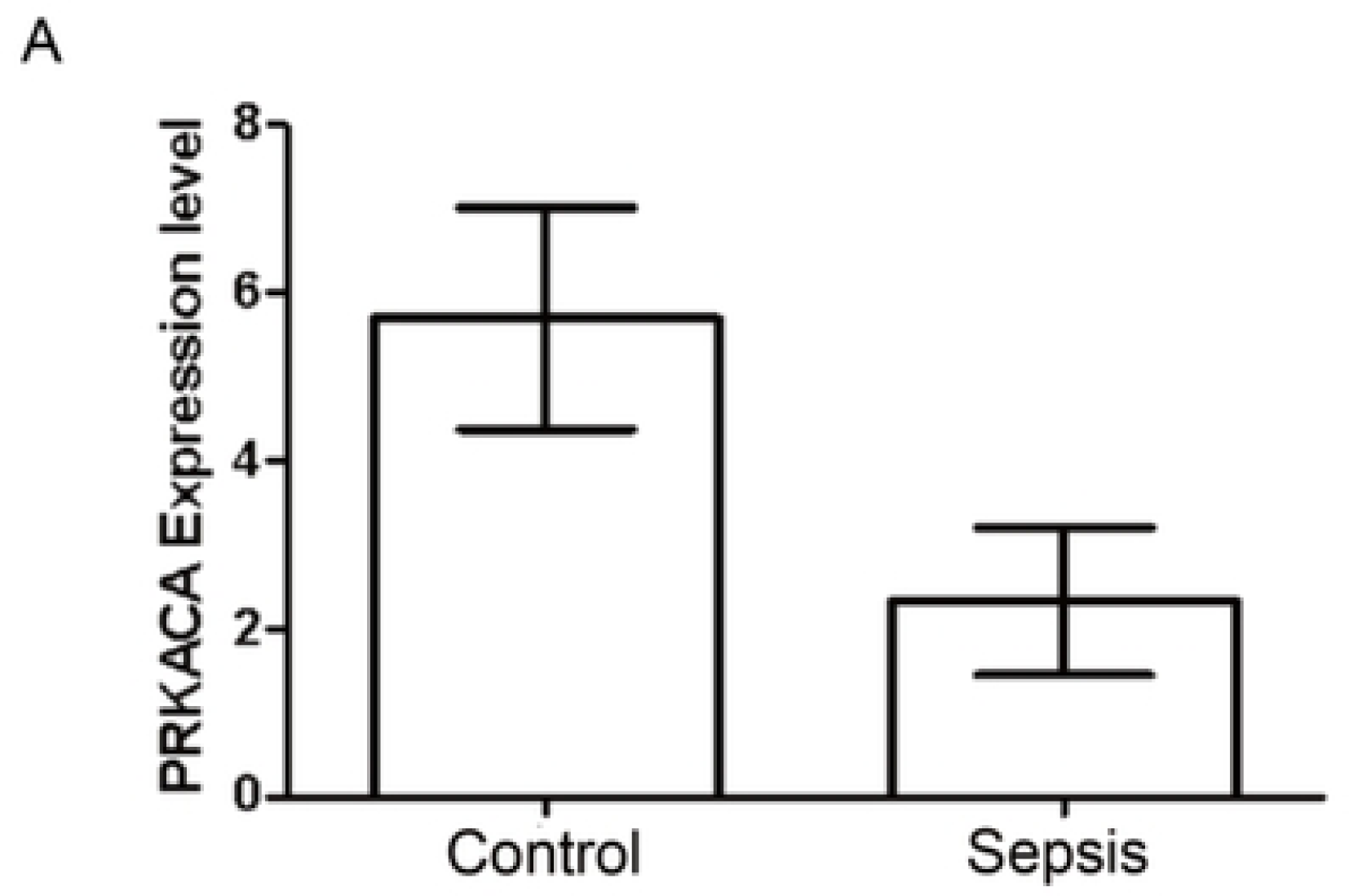
qcpr experiment reveals expression of PRKACA in sepsis and control groups.

**Fig 5B.**
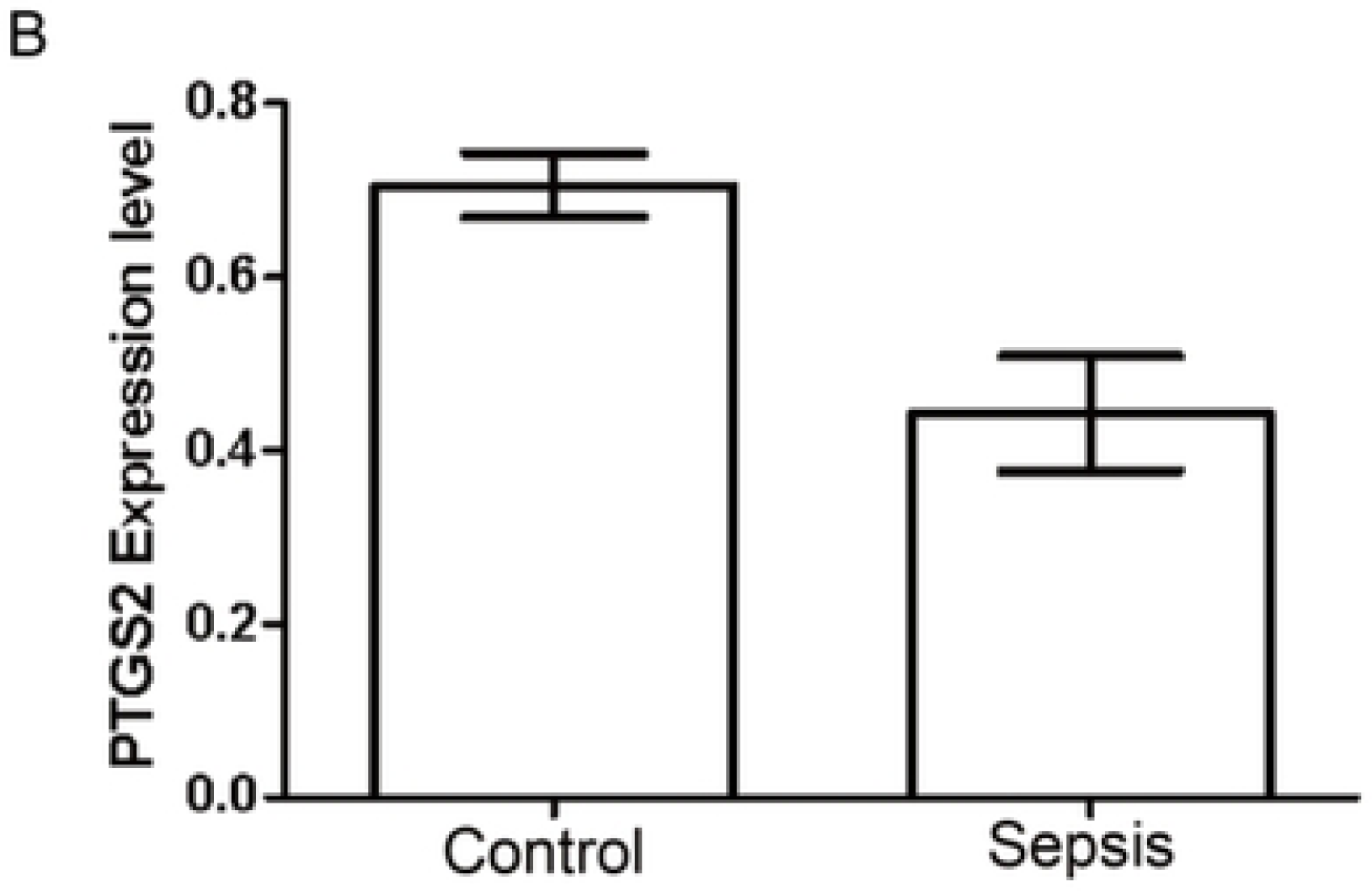
qcpr experiment reveals expression of PTGS2 in sepsis and control groups.

**Fig 5C.**
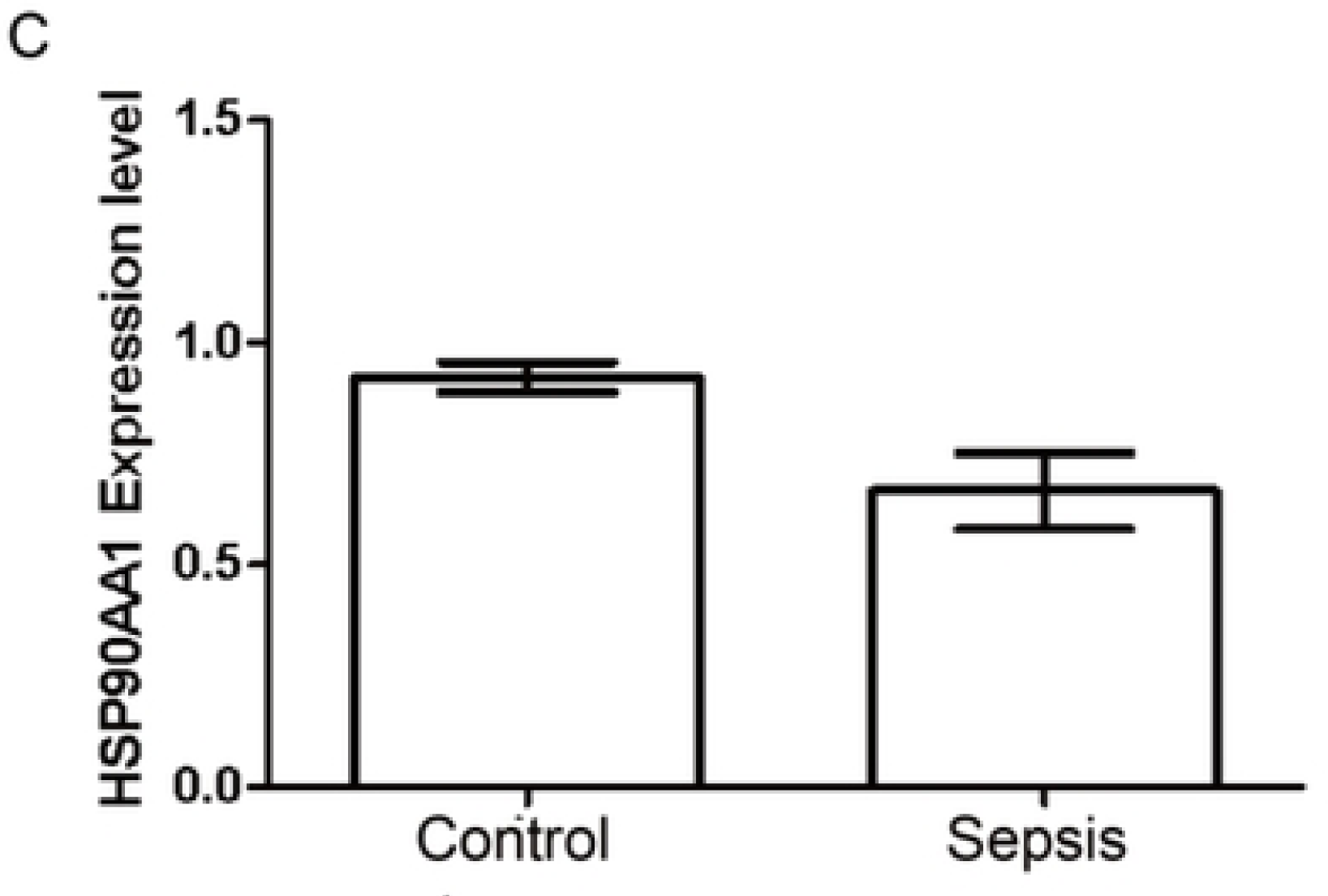
qcpr experiment reveals expression of HSP90AA1 in sepsis and control groups.

**Fig 5D.**
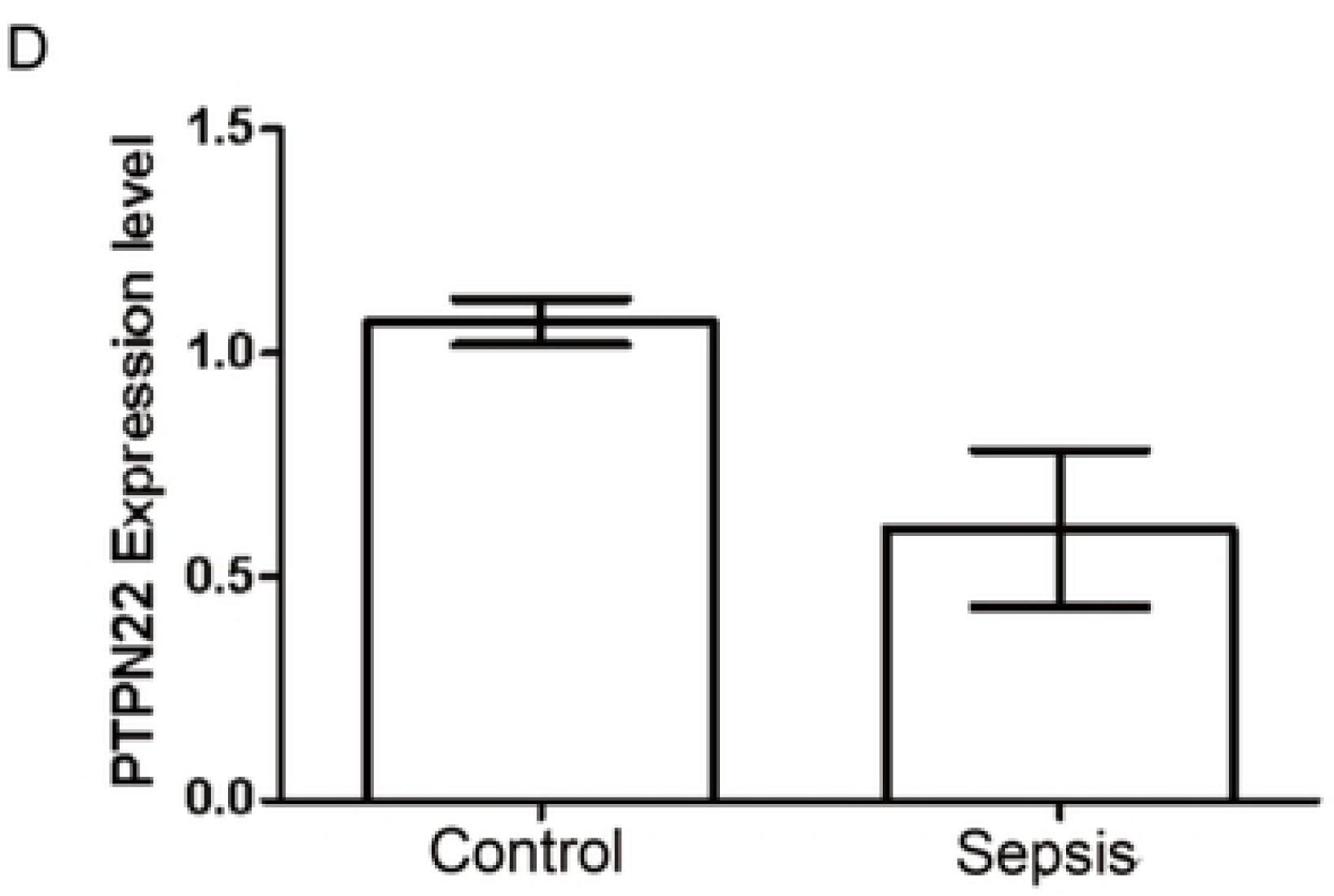
qcpr experiment reveals expression of PTPN22 in sepsis and control groups.

**Fig 5E.**
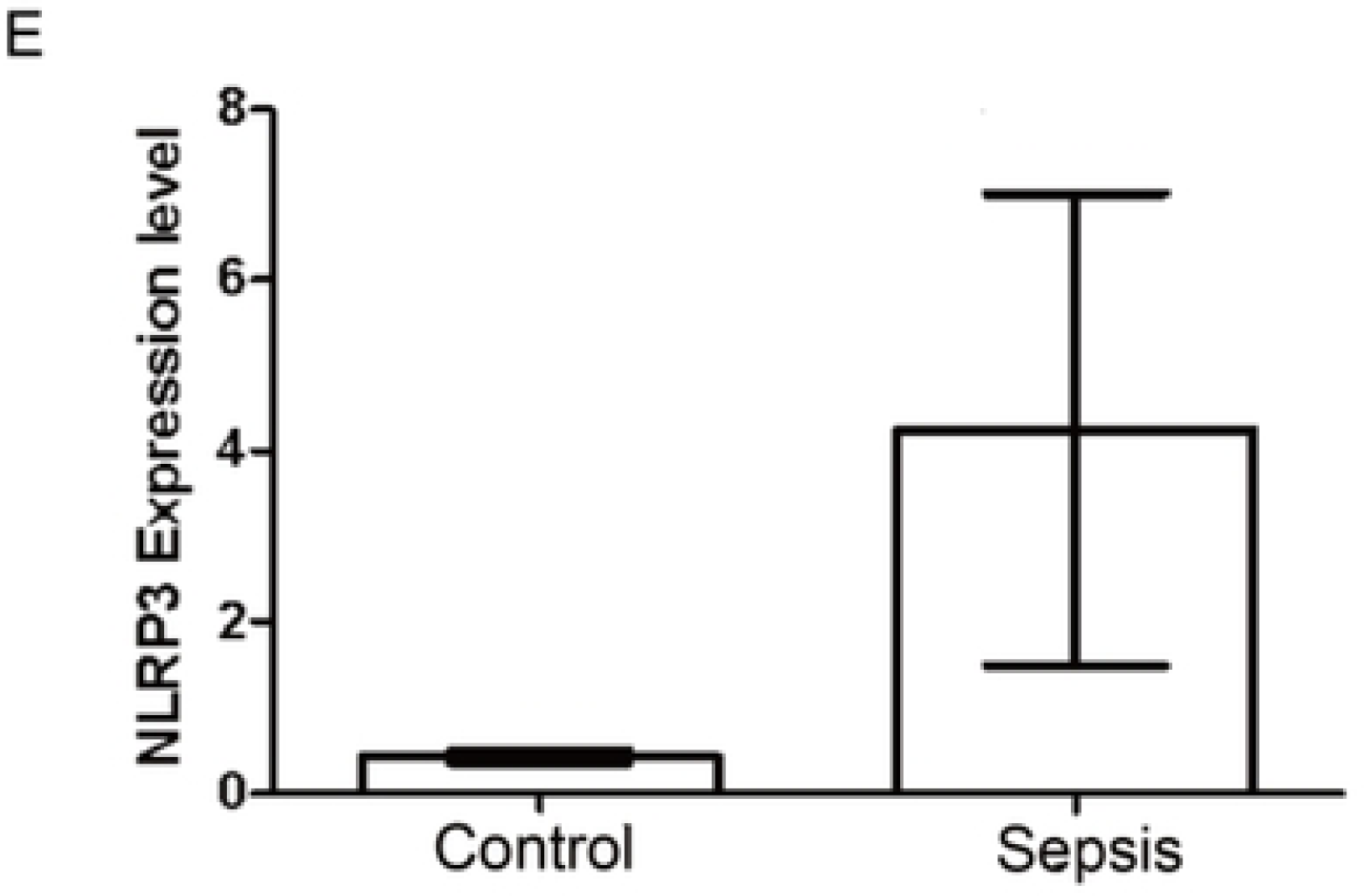
qcpr experiment reveals expression of NLRP3 in sepsis and control groups.

## Discussion

Our study explores the mechanism and process by which palmatine affects pyroptosis in sepsis cells through related signaling pathways. In this study, we identified 9 pyroptosis-related differentially expressed genes (DEGs) between sepsis and control groups and analyzed their correlations. By analyzing the expression of these 9 genes in the dataset and validation set of this study, we found that the upregulation of individual genes (NLRC4) indicated the persistent inflammation of local tissues and the activation of pyroptosis in sepsis, while the downregulation of the remaining genes indicated the low immune function of patients with sepsis pyroptosis, which may be the immunosuppression caused by pyroptosis, reflecting the inflammatory and immune imbalance in patients with sepsis pyroptosis. Previous studies have shown that CXCL8 is closely associated with inflammation and promotes neutrophil recruitment as a chemokine [22]. CXCL8 is a key inflammation-related gene, with connections to multiple other genes, including IL1B, PTGS2, NLRP6, and NLRP1. According to the literature, NLRP1, NLRC4, and NLRP6 are members of the NOD-like receptor (NLR) family. Among these: NLRC4 forms inflammasomes through molecular interactions.

NLRP1 is closely related to the formation of inflammasome. NLRP6 negatively regulates the NF-κB and MAPK signaling pathways [23]. The correlation analysis reveals a strong relationship between NLRP6 and NLRP1, both of which are also linked to CXCL8. Furthermore, NLRC4 is connected to PTGS2 and AKT1, suggesting functional interactions and similarities. NLRP3, another member of the NLR family, is also closely associated with pyroptosis. Recent studies indicate that PTGS2 expression can be induced by inflammatory stimuli [24], playing a critical role in inflammatory diseases. Additionally, PTGS2 is a classic downstream molecule of the NF-κB pathway and interacts with NLRC4 and CXCL8. We also found that IL1B is a well-known proinflammatory cytokine [25]. Our correlation results show that IL1B is closely connected to CXCL8, NLRP1, IRF1, and CASP5. These connections suggest that these inflammation-related factors exhibit synergistic effects and are all intricately linked to cell pyroptosis.

We also performed GO and KEGG enrichment analyses on the 9 pyroptosis-related differentially expressed genes. The results revealed significant enrichment of these genes in the Toll-like receptor signaling pathway, NF-kappa B signaling pathway, NOD-like receptor signaling pathway, and TNF signaling pathway. Among these, the NOD-like receptor signaling pathway serves as the core mechanism of pyroptosis, as NLR family genes directly participate in inflammasome activation. In the NF-kappa B signaling pathway, activation facilitates the release of inflammatory cytokines [26]. Upstream genes such as IRF1 and AKT1 may regulate the expression of NLRP1 and IL1B through NF-κB signaling, thereby amplifying the inflammatory response associated with pyroptosis. Regarding the Toll-like receptor signaling pathway, recent studies have shown that it induces inflammatory responses and triggers antiviral immune defenses [27]. Genes such as PTGS2 and CXCL8 may detect pathogen signals via the TLR pathway, enhancing the inflammatory response in sepsis. These findings further demonstrate that these genes collaboratively participate in the pyroptosis response and play a role in regulating the inflammatory processes in sepsis.

To further investigate the relationship between pyroptosis clusters and immune infiltrating cells, we analyzed the differences in immune infiltration levels between Clusters A and B. The results revealed that the infiltration level of T cells CD4 naïve was higher in Cluster B compared to Cluster A, indicating immune microenvironment activation in Cluster B. Conversely, the infiltration level of plasma cells was lower in Cluster B, suggesting immune microenvironment inhibition. Subsequent GSVA enrichment analysis demonstrated that Cluster A was primarily enriched in immunity and DNA repair, while Cluster B was predominantly enriched in signal transduction and metabolism. From the biological process (BP) analysis, we observed significant enrichment in processes related to immune responses and cytokines. The cellular component (CC) analysis showed significant enrichment in processes related to secretion and intercellular signaling. According to the molecular function (MF) analysis of DEGs between the two clusters, protein binding, nucleotide binding, and metabolism were significantly enriched. KEGG pathway analysis further revealed significant enrichment in inflammatory response-related pathways, immune regulation, and pathogen infection-related pathways. These findings highlight the potential roles of these chemical reactions and biological processes in pyroptosis among sepsis patients.

Overall, these results indicate that the pathways and biological functions enriched in pyroptotic clusters differ significantly among sepsis patients. This underscores the importance of tailoring treatment strategies to specific patient populations based on their pyroptosis-related molecular and immune profiles.

Using network pharmacology, we identified the relevant targets of palmatine and performed GO and KEGG enrichment analyses on these targets. Additionally, by constructing a pathway network linking these targets to KEGG pathways, we determined the biological functions and effects that palmatine may exert through associated signaling pathways. KEGG enrichment analysis revealed that the pathways enriched in these targets are primarily involved in anti-inflammatory and immune regulation, hormone regulation, metabolic regulation, and cell signaling. Notably, PTGS2 was enriched in multiple inflammatory and signaling pathways, such as the IL-17 signaling pathway and NF-kappa B signaling pathway. Furthermore, both PTGS1 and PTGS2 were enriched in the Arachidonic acid metabolism pathway, suggesting their synergistic roles in lipid metabolism and inflammatory responses. When analyzing the genes and signaling pathways associated with sepsis pyroptosis alongside the signaling pathways linked to palmatine targets, we identified pathways with overlapping or similar functions. These include the IL-17 signaling pathway, NF-kappa B signaling pathway, NOD-like receptor signaling pathway, and TNF signaling pathway. These pathways are strongly associated with PTPN22 and NLRP3, underscoring their potential roles in the pyroptosis response and inflammation in sepsis. Other studies have shown that PTPN22 can interact with and dephosphorylate NLRP3[28] and PTPN22 usually regulates the release of pro-inflammatory cytokines by inhibiting NF-κB signaling pathway. We noticed that expression level of PTPN22 in sepsis patients was downregulated compared with control group, indicating in sepsis, normal regulation of inflammation was disrupted. Meanwhile, expression level of NLRP3 in sepsis patients was upregulated compared with control group, indicating increasing inflammatory response and pyroptosis.

We verified the expression of genes active in the target of palmatine and genes closely related to the above signaling pathways in sepsis and control groups by qPCR experiments. The results showed that PRKACA, PTGS2, HSP90AA1 and PTPN22 were all downregulated, indicating that palmatine may be associated with anti-inflammatory and immunomodulatory effects through signaling pathways. Besides, NLRP3 was upregulated. In addition, since these genes are direct targets of palmatine, palmatine can also be used as a drug for sepsis. Among them, PTGS2 not only plays a role in sepsis pyroptosis, but is also the target of palmatine, which also shows that palmatine can act on sepsis pyroptosis.

We investigated the signaling pathways associated with pyroptosis-related genes in sepsis patients, classified these patients into subtypes based on these genes, and conducted analyses such as immune infiltration and GSVA enrichment to explore the biological differences between the subtypes. This work lays the groundwork for subsequent treatment strategies and further research. Additionally, through network pharmacology, we identified the relevant targets of palmatine and the KEGG signaling pathways associated with these targets. We observed overlapping functions and similarities between the KEGG signaling pathways related to palmatine targets and those related to sepsis pyroptosis genes. Notably, PTGS2 was found to play a critical role not only in palmatine’s mechanism of action but also in sepsis pyroptosis, providing further evidence that palmatine may have a therapeutic effect on patients with sepsis pyroptosis. Furthermore, pathways associated with NLRP3 and PTPN22 were also confirmed in the genes of sepsis pyroptosis patients, elucidating a complete gene-pathway mechanism of action.

Despite these findings, our study has some limitations. First, we lacked detailed information on the clinical characteristics of the patients analyzed. Second, additional wet-lab experiments are needed to validate the effects of palmatine’s targets and components on sepsis-related pyroptosis in patients.

## Conclusion

In this study, we explored the genes and signaling pathways associated with pyroptosis in sepsis and classified sepsis into distinct clusters based on these genes. This work lays a foundation for understanding the impact of pyroptosis-related genes on sepsis classification. Furthermore, we investigated the targets and signaling pathways of palmatine, shedding light on its role in modulating pyroptosis in sepsis cells.

## Availability of data and materials

The datasets generated during the current study are available in the Network_ Pharmacology repository: TCMSP database, https://www.tcmsp-e.com/#/databaseacology.

## Author contributions

GuangzhaoYan and Yanyan Wangdesigned the study, analyzed the data, and wrote and organized the paper.All authors have read the final paper and approved it for publication.

## Ethics approval and consent to participate

This study was approved by the Ethics Committee of Zhejiang Provincial People’s Hospital(KT2025002) 。 and was performed in accordance with the Declaration of Helsinki for medical research involving human subjects. Written informed consent was acquired from each participant

## Supplementary manerials

You can see KEGG id and corresponding name in supplementary materials.

## Reference

1. Cecconi M, Evans L, Levy M, Rhodes A. Sepsis and septic shock. The Lancet. 2018 Jul 7;392(10141):75–87. doi:10.1016/S0140-6736(18)30696-2

2. Joffre J, Hellman J, Ince C, Ait-Oufella H. Endothelial Responses in Sepsis. American Journal of Respiratory and Critical Care Medicine. 2020 Aug 1;202(3):361–70. doi:10.1164/rccm.201910-1911TR

3. Cecconi M, Evans L, Levy M, Rhodes A. Sepsis and septic shock. The Lancet. 2018 Jul 7;392(10141):75–87. doi:10.1016/S0140-6736(18)30696-2

4. Xie J, Wang H, Kang Y, Zhou L, Liu Z, Qin B, et al. The Epidemiology of Sepsis in Chinese ICUs: A National Cross-Sectional Survey. Critical Care Medicine. 2020;48(3):e209–18. doi:10.1097/CCM.0000000000004155

5. Li Y, Sheng H, Qian J, Wang Y. The Chinese Medicine Babaodan Suppresses LPS-Induced Sepsis by Inhibiting NLRP3-Mediated Inflammasome Activation. Journal of ethnopharmacology. 2022 Jun 28;292:115205. doi:10.1016/j.jep.2022.115205

6. Long J, Song J, Zhong L, Liao Y, Liu L, Li X. Palmatine: A review of its pharmacology, toxicity and pharmacokinetics. Biochimie. 2019;162:176–84. doi:10.1016/j.biochi.2019.04.008

7. Ekeuku SO, Pang KL, Chin KY. Palmatine as an Agent Against Metabolic Syndrome and Its Related Complications: A Review. Drug Des Devel Ther. 2020 Nov 17;14:4963–4974. doi:10.2147/DDDT.S280520

8. Dickson K, Lehmann C. Inflammatory Response to Different Toxins in Experimental Sepsis Models. Int J Mol Sci. 2019 Sep 5;20(18):4341. doi:10.3390/ijms20184341

9. Chen G, Wang X, Liu C, Zhang M, Han X, Xu Y. The interaction of MD-2 with small molecules in huanglian jiedu decoction play a critical role in the treatment of sepsis. Frontiers in Pharmacology. 2022 Sep 9;13:947095. doi:10.3389/fphar.2022.947095

10. Chen G, Xu Y, Jing J, Mackie B, Zheng X, Zhang X, et al. The anti-sepsis activity of the components of Huanglian Jiedu Decoction with high lipid A-binding affinity. International Immunopharmacology. 2017;46:87–96. doi:10.1016/j.intimp.2017.02.025

11. Yan B, Wang D, Dong S, Cheng Z, Na L, Sang M, et al. Palmatine inhibits TRIF-dependent NF-κB pathway against inflammation induced by LPS in goat endometrial epithelial cells. International Immunopharmacology. 2017;45:194–200. doi:10.1016/j.intimp.2017.02.004

12. Tang C, Hong J, Hu C, Huang C, Gao J, Huang J, et al. Palmatine Protects against Cerebral Ischemia/Reperfusion Injury by Activation of the AMPK/Nrf2 Pathway. Oxidative Medicine and Cellular Longevity. 2021 Mar 11;2021:6660193. doi:10.1155/2021/6660193

13. Kovacs SB, Miao EA. Gasdermins: Effectors of Pyroptosis. Trends in Cell Biology. 2017;27(9):673–84. doi:10.1016/j.tcb.2017.05.005

14. Li W, Sun J, Zhou X, Lu Y, Cui W, Miao L. Mini-Review: GSDME-Mediated Pyroptosis in Diabetic Nephropathy. Frontiers in Pharmacology. 2021 Nov 16;12. doi:10.3389/fphar.2021.780790

15. Broz P, Pelegrín P, Shao F. The gasdermins, a protein family executing cell death and inflammation. Nature Reviews Immunology. 2019 Nov 5;20(3):143–57. doi:10.1038/s41577-019-0228-2

16. Li S, Feng L, Li G, Liu R, Ma C, Wang L, et al. GSDME-dependent pyroptosis signaling pathway in diabetic nephropathy. Cell Death Discovery. 2023 May 11;9(1). doi:10.1038/s41420-023-01452-8

17. Ding W, Huang L, Wu Y, Su J, He L, Tang Z, et al. The role of pyroptosis-related genes in the diagnosis and subclassification of sepsis. PLOS ONE. 2023 Nov 8;18(11):e0293537. doi:10.1371/journal.pone.0293537

18. Gou X, Xu D, Li F, Hou K, Fang W, Li Y. Pyroptosis in stroke-new insights into disease mechanisms and therapeutic strategies. Journal of Physiology and Biochemistry. 2021 May 4;77(4):511–29. doi:10.1007/s13105-021-00817-w

19. Aglietti RA, Dueber EC. Recent Insights into the Molecular Mechanisms Underlying Pyroptosis and Gasdermin Family Functions. Trends in Immunology. 2017;38(4):261–71. doi:10.1016/j.it.2017.01.003

20. Zheng X, Chen W, Gong F, Chen Y, Chen E. The Role and Mechanism of Pyroptosis and Potential Therapeutic Targets in Sepsis: A Review. Frontiers in Immunology. 2021 Jul 7;12:711939. doi:10.3389/fimmu.2021.711939

21. Liang S, Xing M, Chen X, Peng J, Song Z, Zou W. Predicting the prognosis in patients with sepsis by a pyroptosis-related gene signature. Frontiers in Immunology. 2022 Dec 21;13:1110602. doi:10.3389/fimmu.2022.1110602

22. Hou Y, Huttenlocher A. Advancing chemokine research: the molecular function of CXCL8. Journal of Clinical Investigation. 2024 May 15;134(10):e180984. doi:10.1172/JCI180984

23. Zhou Y, Yu S, Zhang W. NOD-like Receptor Signaling Pathway in Gastrointestinal Inflammatory Diseases and Cancers. International Journal of Molecular Sciences. 2023 Sep 25;24(19):14511. doi:10.3390/ijms241914511

24. Tanaka M, Shirakura K, Takayama Y, Μatsui M, Watanabe Y, Yamamoto T, et al. Endothelial ROBO4 suppresses PTGS2/COX-2 expression and inflammatory diseases. Communications Biology. 2024 May 18;7(1). doi:10.1038/s42003-024-06317-z

25. Rawat M, Nighot M, Al-Sadi R, Gupta Y, Viszwapriya D, Yochum G, et al. IL1B Increases Intestinal Tight Junction Permeability by Up-regulation of MIR200C-3p, Which Degrades Occludin mRNA. Gastroenterology. 2020;159(4):1375–89. doi:10.1053/j.gastro.2020.06.038

26. Kang C, Li X, Liu P, Liu Y, Niu Y, Zeng X, et al. Tolerogenic dendritic cells and TLR4/IRAK4/NF-κB signaling pathway in allergic rhinitis. Frontiers in Immunology. 2023 Oct 17;14:1276512. doi:10.3389/fimmu.2023.1276512

27. Duan T, Du Y, Xing C, Wang HY, Wang RF. Toll-Like Receptor Signaling and Its Role in Cell-Mediated Immunity. Frontiers in Immunology. 2022 Mar 3;13:812774. doi:10.3389/fimmu.2022.812774

28. Spalinger MR, Lang S, Gottier C, Dai X, Rawlings DJ, Chan AC, et al. PTPN22 regulates NLRP3-mediated IL1B secretion in an autophagy-dependent manner. Autophagy. 2017 Aug 25;13(9):1590–601. doi:10.1080/15548627.2017.1341453

